# Tissue-specific adaptation of human T cells is preserved during tissue inflammation

**DOI:** 10.64898/2026.04.09.717255

**Authors:** Nicole B. Potchen, Hugh R. MacMillan, Eva Domenjo-Vila, Andrew J. Konecny, Alexis K. Taber, Caitlin S. DeJong, Geervani Daggupati, Swati Shree, Stephen A. McCartney, Shelton W. Wright, Evan W. Newell, Douglas R. Dixon, Martin Prlic

## Abstract

T cells play an essential role in protecting tissues against pathogens and regulating tissue homeostasis. Previous studies highlight that T cells display tissue-specific phenotypic and functional properties, suggesting that T cells adapt to their local environment. Whether inflammation disrupts tissue-specific T cell adaptation remains poorly understood. To address this open question, we examined the T cell compartment – including conventional CD4 and CD8 T cells, regulatory T cells, γδ T cells, and MAIT cells – from healthy and inflamed human mucosal tissues. Using high-parameter spectral flow cytometry, we examined phenotype *ex vivo* and the functional capacity following stimulation, utilizing conventional gating and unsupervised clustering analysis approaches. Overall, we analyzed 65 tissue samples including mild, moderate, and severely inflamed oral gingiva, healthy and inflamed lung, along with healthy and inflamed tissue from the decidual-placental interface. Across these mucosal barrier tissues, we find that tissue location plays a dominant role in shaping the composition, phenotype, and functional capacity of the T cell compartment. Importantly, these tissue-specific adaptations were largely maintained during states of tissue inflammation. This included the ability to exert tissue repair functions, which was preserved across T cell subsets, even in severely inflamed tissues.

## Introduction

Human T cells have distinct phenotypic and functional properties across different tissues and organs. This was initially demonstrated by Farber and colleagues who examined the phenotype and functional properties of conventional CD4 and CD8 T cells isolated from secondary lymphoid tissues, lung, jejunum, ileum and colon of trauma-induced, deceased organ donors^1^. This study furthermore established CD69 expression as a biomarker for tissue-residence for human CD4 T cells, and CD69 and CD103 co-expression to indicate tissue-residence for CD8 T cells^1^. Follow-up studies highlighted that expression of CD69 by human CD4 and CD8 T cells is associated with a core transcriptional signature akin to bona fide murine tissue-resident T cells^2^. Human tissue-resident memory CD8 T cells (T_RM_) show tissue-specific adaptation such as distinct cytotoxic properties across skin, lung, and jejunum, and are also clonally distinct across these tissue sites^3–7^. A recent study indicated that within the intestine, distinct tissue niches drive unique phenotypic and functional properties in CD8 T cells, thus providing further compelling evidence that T_RM_ tissue location and functional state are fundamentally linked^8^. Similarly, conventional CD4 T cells (CD4 Tconv) and regulatory T cells (CD4 Treg) also have distinct phenotypic and functional properties across tissues^9–12^. Further evidence points to innate-like T cells also adapting in a tissue-specific manner. Human γδ T cells differ by tissue location with cells from adult lung, spleen, lymph nodes (LNs), and gut tissue clustering differentially by tissue based on their expression of transcripts associated with effector and repair functions^13^. Finally, human mucosal-associated invariant T (MAIT) cells across tissues have distinct T cell receptor (TCR) repertoires and cytokine expression profiles^14,15^.

Whether tissue-specific adaptations are maintained during tissue inflammation is unclear. Inflammation involves coordinated immune cell recruitment and activation and thus has the potential to profoundly alter the T cell compartment within tissue^16^. Memory phenotype CD8 T cells can be activated by pro-inflammatory signals alone; this event has been coined bystander-activation or inflammation-driven activation of T cells^17,18^. Inflammation-driven activation of memory CD8 T cells leads to increased expression of CD69, PD-1, and TOX, along with expression of IFNγ and Granzyme B (GzmB)^19–23^ which can have protective properties^24,25^ but can also lead to tissue pathologies^26^. Such bystander-activated CD8 T cell-mediated pathology has been demonstrated in a human cohort following infection with Hepatitis A virus (HAV)^27^. For CD4 T cells, inflammation and associated tissue pathology is often associated with expression of IL-17, but it is noteworthy that IL-17 should not be simply viewed as a pro-inflammatory cytokine^28^ or indicator of excessive inflammation^29^. Importantly, in addition to mediating tissue protection and pathology, T cells also play a role in restoring tissue homeostasis via secretion of IL-22 and the epidermal growth factor receptor (EGFR)-ligand amphiregulin (AREG)^30–32^. Tregs express AREG in response to IL-18 and IL-33^33^, while a recent study demonstrates that human memory CD8 T cells express AREG in response to TCR stimulation^34^.

While there has been substantial progress in our understanding of the T cell compartment and its tissue-specific functional properties in healthy human tissues, the extent to which inflammation alters these functional properties remains an open question. We sought to address if inflammation overrides any tissue-specific T cell adaption in human tissues. Specifically, we aimed to determine if inflammation alters the composition of the T cell compartment and associated functional properties, and if it affects the ability of T cells to maintain their tissue-regenerative functions. To accomplish this, we collected healthy and inflamed tissues from 3 distinct mucosal barrier sites: oral mucosa (gingiva), lung, and the decidual-placental interface (DPI). We developed a high-parameter flow cytometry panel containing 37 markers to define the composition, phenotype, and functional capacity of the T cell compartment: CD4 Tconv, CD4 Treg, CD8 CD103+ T_RM_ and CD8 CD103- circulating T cells, along with the innate-like MAIT and γδ T cell subsets. Overall, we found that the composition, phenotype and function of the T cell compartment was driven more by tissue location than inflammation, indicating that T cells largely maintain their tissue-specific adaptations during inflammation. Our data emphasize the need to assess and consider inflammation in a tissue-specific manner. Furthermore, our data highlight that T cells maintain their tissue-regenerative capacity even in severely inflamed tissues, thus pinpointing T cells as a therapeutic target to treat inflammatory diseases.

## Results

### The inflammation state in the oral mucosa has a modest effect on composition and phenotype of the T cell compartment

To determine the impact of inflammation on the mucosal T cell compartment, we first examined phenotypic differences across mildly, moderately, and severely inflamed gingival tissues. These tissues are discarded from routine surgeries and represent a wide range of inflammatory states. We assigned clinically healthy gingival tissue and tissue with mild gingivitis as “mild inflammation” since even clinically healthy tissue contains a baseline level of inflammation, which may allow for constant immune surveillance^35^. Next, we assigned tissues with pronounced gingivitis transitioning to periodontitis as “moderate inflammation” and those with periodontitis as “severe inflammation”. While gingivitis is reversible with proper dental hygiene, periodontitis is an irreversible, chronic disease state without clinical intervention^36^. A full sample list with clinical information can be found in Supplemental Table 1. A 37-color spectral flow cytometry panel was utilized to examine phenotype (no stimulation) and functional capacity (following stimulation) of the T cells isolated from gingival tissues (***Figure 1A****, panel and gating; Supplemental Figure 1-2*). As a technical control, a PBMC sample from a single leukapheresis donor was included in every flow cytometry experiment (*Supplemental Figure 3*).

**Figure 1:**
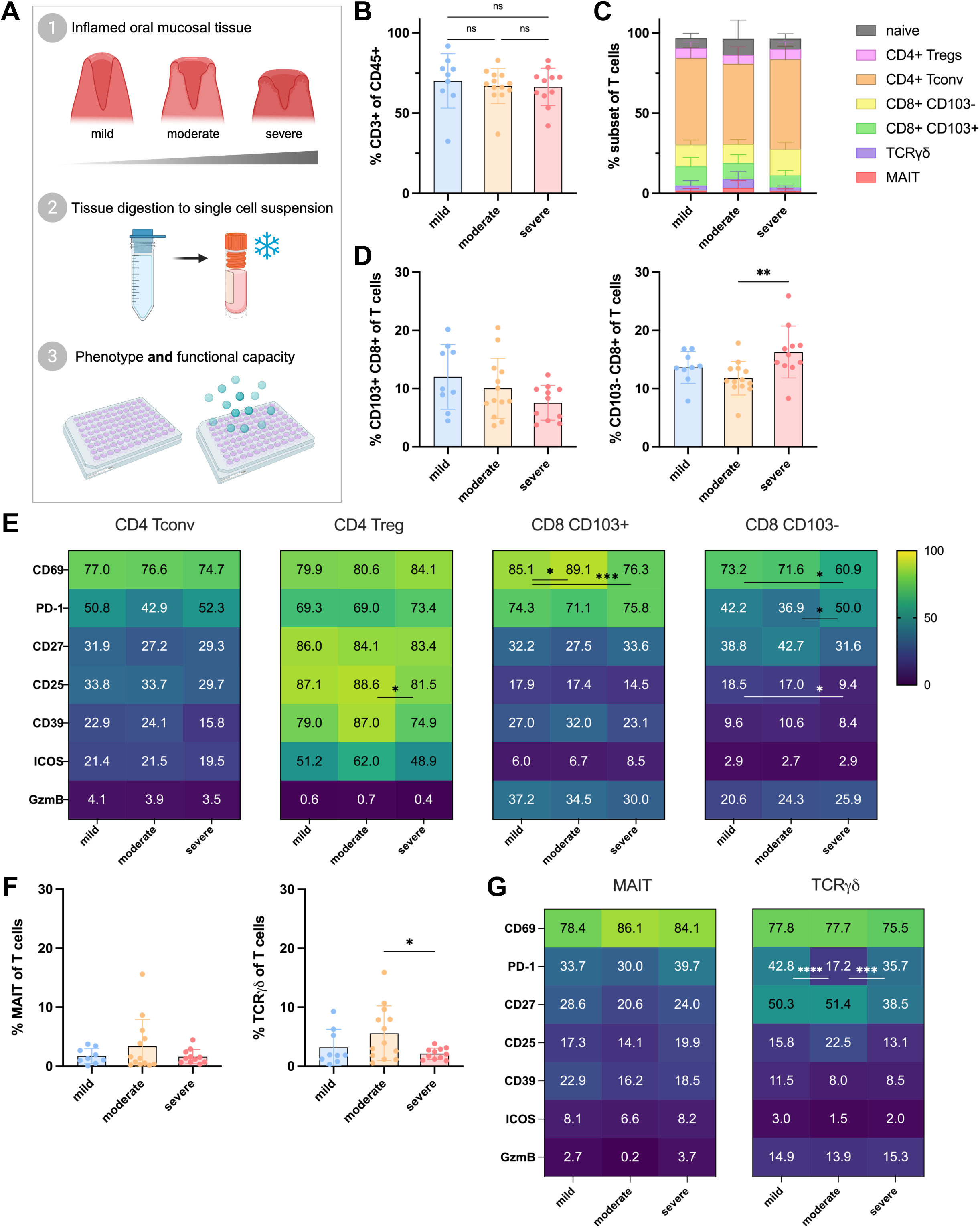
Inflammation in the oral mucosa modestly affects composition and phenotype of the T cell compartment. **A.** Experimental layout, described in detail in methods. Briefly, oral mucosa tissue of varying inflammation states was sent from the clinic to lab, kept at 4°C. Tissue was digested mechanically and enzymatically prior to storage of cryovials with single cell suspension in liquid nitrogen. Upon experiment, samples were thawed and assessed directly ex vivo (Figures 1-2) or following a 6h PMA/I stimulation at 37°C (Figure 3). **B.** Frequency of CD3+ T cells of CD45+ cells across inflammation state, pre-gated on lymphocytes, single cells, and live (*Supplementary* Figure 1). **C.** Of CD3+ CD45+ cells, frequency of each T cell subset. **D.** Frequency of subset (left to right, CD8+ CD103+ resident, and CD8+ CD103- circulating) of CD3+ CD45+ T cells across inflammation state. For **B-D.** T cell subsetting, n = 9 mild, 13 moderate, 11 severely inflamed oral mucosa tissues. **E.** For each conventional subset, heatmaps depict frequency of activation markers across inflammation status. Phenotyping of T cell subsets must meet minimum 25 cell threshold; sample size is consistent with above unless noted; n = 8 mild, 11 moderate for Tregs. Tconv, CD103+ and CD103- all met minimum threshold. **F.** Frequency of MAIT cells and γδ T cells of CD3+ CD45+ T cells. **G.** For innate-like T cells, frequency of activation markers across inflammation. Minimum threshold of 25 cells; n = 7 mild, 9 moderate for MAIT cells; n = 8 mild for γδ T cells. For all plots, ns (not significant), *p < 0.05, ***p < 0.001, ****p < 0.0001 by ordinary one-way ANOVA with Tukey’s multiple comparison test.

We first examined the frequency of T cells (CD3+ cells) within the CD45+ cell compartment and found no changes between mild, moderate, and severely inflamed gingival tissue (***Figure 1B***). Of note, a small population of naïve T cells (CCR7+ CD45RA+) was present in each tissue sample. These naïve T cells presumably stem from the blood that is inherently present in the tissue vasculature. We excluded these naïve T cells from our manual gating analysis approach. We further analyzed the T cell compartment by first defining innate-like T cell subsets within the CD3+ population, followed by subsetting the remaining CD4 and CD8 T cells into conventional CD4 T cells (CD4 Tconv), regulatory CD4 T cells (Treg), CD103+ (tissue-resident phenotype) CD8 T cells and CD103- (circulating) CD8 T cells (*Supplemental Figure 1B*). We then assessed if T cell subset composition is altered with inflammation. We observed stable frequencies of most T cell subsets, including CD4 Tconv, CD4 Treg, and CD103+ CD8 T cells (***Figure 1C***). We noticed a slight reduction in CD103+ CD8 T cells within the total T cell population (p = 0.1026), but a significant increase in circulating CD103- CD8 T cells in severely inflamed tissue compared to mild inflammation (***Figure 1D***).

We next examined if the expression of activation, co-stimulation, and exhaustion markers change with inflammation severity. To visualize similarities and differences in expression of multiple proteins across different tissues, we chose a heatmap-based approach to displaying expression frequency. For each protein, the mean expression frequency is shown on all heatmaps, and the median and data range for each protein are shown in *Supplemental Table 2.* Within conventional CD4 and CD8 T cell subsets, we observed only few changes between mild, moderate, and severely inflamed gingiva (***Figure 1E***). Pro-inflammatory cytokines can increase the expression of CD69, PD-1, and GzmB by memory CD8 T cells ^20^, but we did not observe an inflammation-associated increase of these proteins (***Figure 1E***). On the contrary, we noted reduced expression of CD69 in both the CD8 CD103+ and CD103- populations in severely inflamed compared to mildly inflamed tissues. In the CD4 Tconv population, there were no differences in these phenotypic markers. In general, we observed relative phenotypic stability across inflammation status within each T cell subset (*Supplemental Figure 4*).

Next, we examined innate-like T cell subsets including MAIT and γδ T cells, which can also be activated by pro-inflammatory stimuli^37,38^. While there were no statistically significant differences in MAIT cell frequencies and phenotypic markers across inflammation status, we found a significant reduction in γδ T cells in severely inflamed compared to moderately inflamed tissue (***Figure 1F***). Within the moderately inflamed condition, we also observed a pronounced reduction of PD-1 expression in the γδ T cell population (***Figure 1G***). The concomitant increase in the abundance of γδ T cells (as a % of CD3+ T cells) and decreased PD-1 expression could potentially be due to recruitment of γδ T cells from the blood to moderately inflamed tissue. While we observed some phenotypic differences in our dataset as tissue became increasingly inflamed, we noted that the T cell profile within all oral mucosal tissues remained overall rather stable even in severely inflamed tissues. We next considered that a smaller proportion of T cells may respond in a TCR-dependent manner.

### Biomarkers of TCR-mediated activation do not significantly change across inflammation states in the oral mucosa

We next assessed biomarker pairs indicative of TCR-mediated activation^39–41^. We examined co-expression of CD39+ PD-1+ (for CD8+ cells^40,41^) as well as ICOS+ PD-1+ (for CD4+ cells^39^) across mild, moderate, and severely inflamed oral mucosal tissues. Across gingiva of varying inflammation states, there were no differences in frequency of expression of each of these populations. CD39, PD-1 and ICOS are also expressed on highly activated Tregs, and we consistently observed the highest frequency of expression in CD4 Tregs (***Figure 2A-B***). In line with this observation that TCR-mediated activation did not change across inflammation states, we also observed no changes in Ki67 expression, an indicator of recent cell proliferation, across inflammation status (***Figure 2C-D***). Lastly, we examined TIM3 and PD-1 expression, a biomarker pair indicative of T cell exhaustion. We observed unaltered co-expression of these markers regardless of inflammation state (***Figures 2C-D***). Together, these data indicate that biomarkers of T cell activation and proliferation remained stable across a range of inflammatory disease states within the gingiva. Overall, we did not detect evidence of inflammation- or TCR-driven activation of the T cell compartment in the inflamed gingiva. To assess the ability of the T cells to respond to activation we next interrogated T cell functional properties.

**Figure 2:**
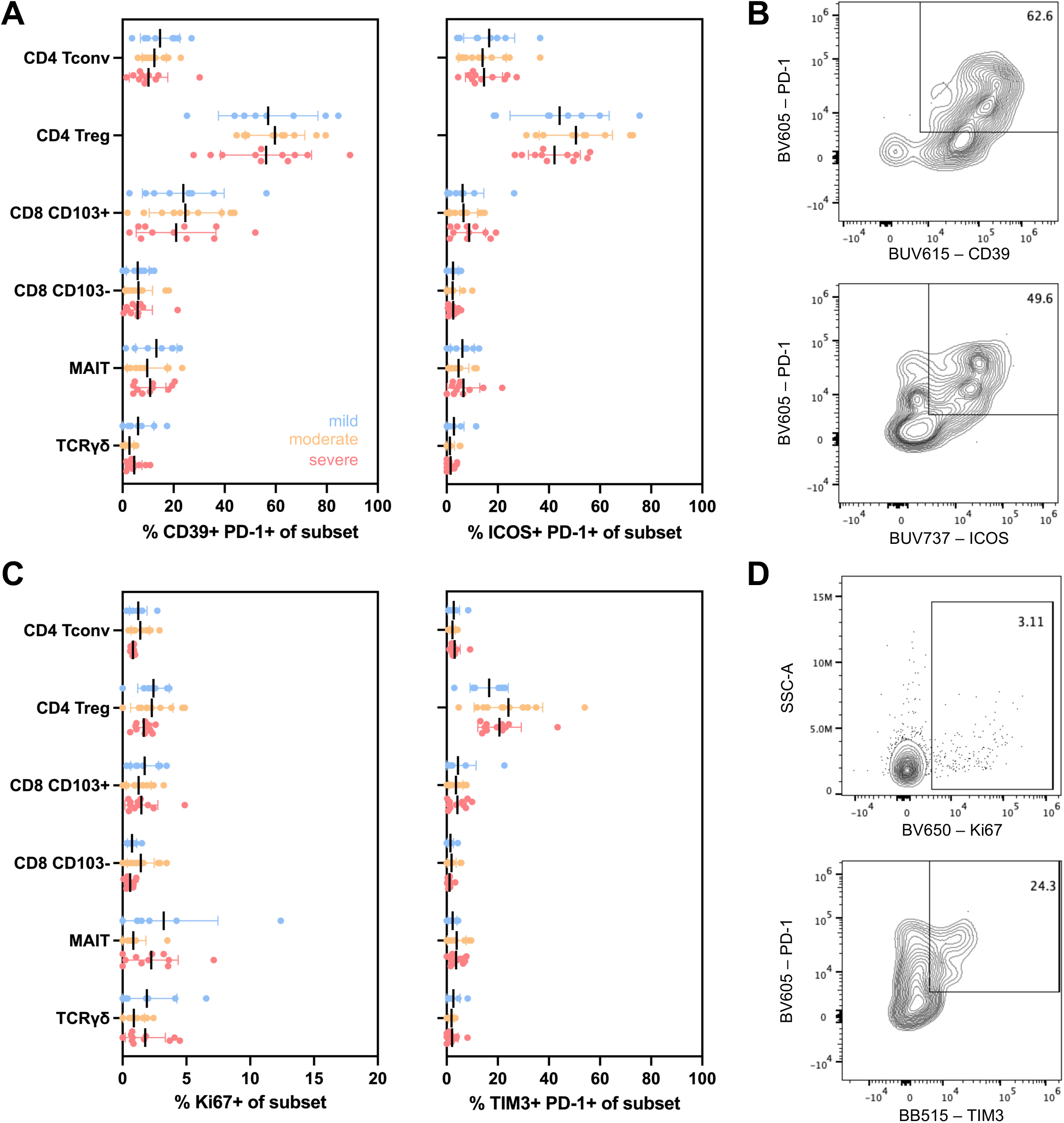
Biomarkers of T cell receptor (TCR)-mediated activation do not significantly change across inflammation states in the oral mucosa. **A.** Expression of CD39+ PD-1+ (right) and ICOS+ PD-1+ (left) among all T cell subsets across inflammation (blue, mild; yellow, moderate; red, severely inflamed). No statistical differences within each subset across inflammation. **B.** Representative flow cytometry plots of CD39 (BUV615) by PD-1 (BV605) expression (top) and ICOS (BUV737) by PD-1 (BV605) expression (bottom) within CD4 Tconv cells isolated from severely inflamed oral mucosal tissue. **C.** From left to right, expression of Ki67+ and TIM3+ PD-1+ across T cell subsets across inflammation status. No statistical differences within each subset. **D.** Representative flow cytometry plots of Ki67 (BV650) expression (top) and TIM3 (BB515) by PD-1 (BV605) expression (bottom) within CD4 Tconv cells isolated from moderately inflamed (top) or severely inflamed (bottom) oral mucosal tissue. No significant differences by ordinary one-way ANOVA with Tukey’s multiple comparison test.

### Functional capacity for T cell-mediated tissue healing and cytokine production is maintained in inflamed oral mucosa

To examine if the functional capacity of T cell subsets varies across inflammation, we stimulated T cells isolated from gingival tissue for 6 hours with PMA/ionomycin (***Figure 1A***). We define the functional capacity simply as the ability of a T cell to express protein following this brief, yet broad, activation stimulus. Within CD4 and CD8 subsets, we observed no statistically significant differences in AREG, IFNγ, TNFα, IL-17A, IL-17F, IL-2, or IL-22 expression across inflammation states (***Figure 3A****, Supplemental Figure 5*). Even the distribution of datapoints for each T cell subset was similar across mildly, moderately, and severely inflamed gingiva (***Figure 3B***).

**Figure 3:**
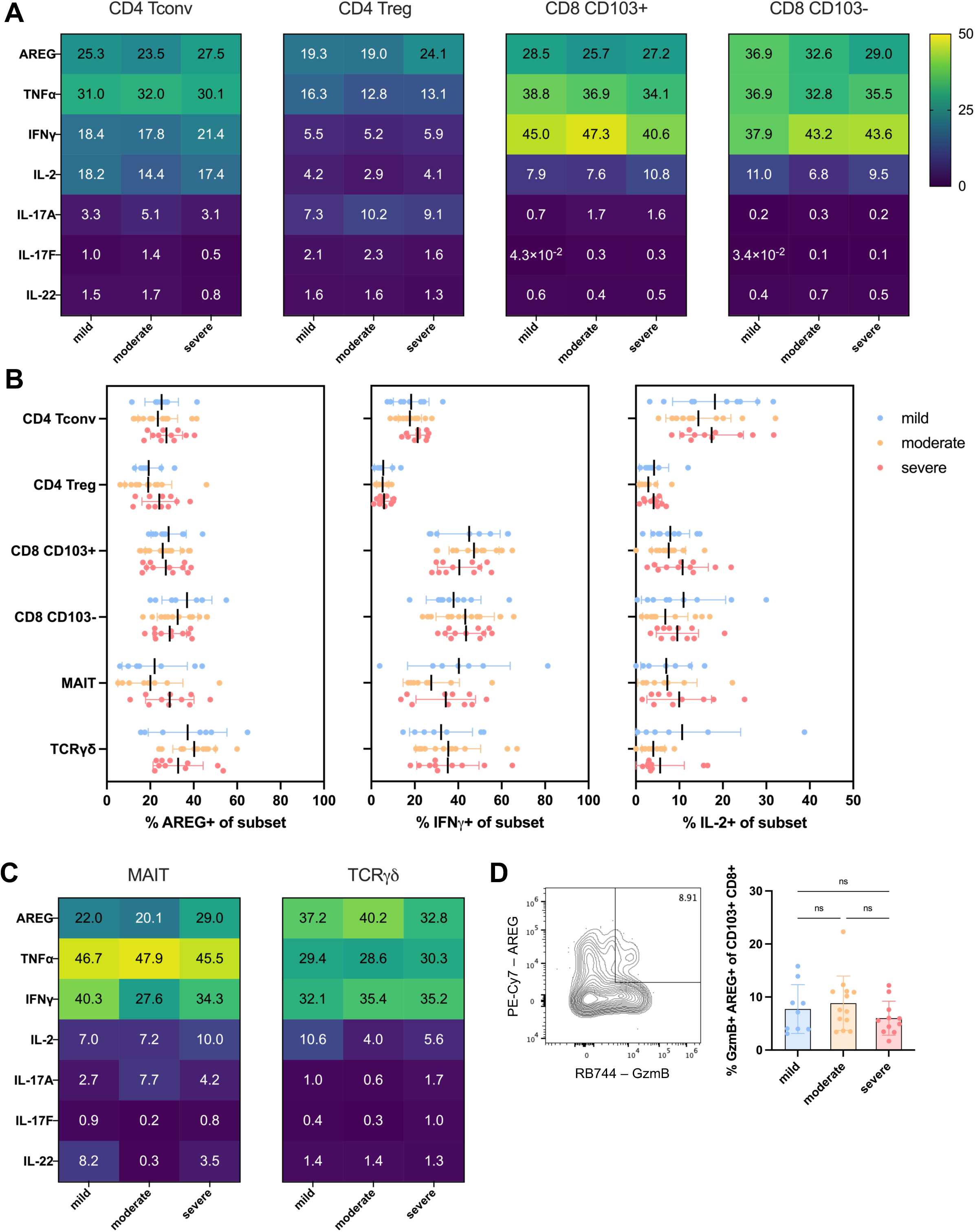
Functional capacity of T cell subsets is maintained in inflamed oral mucosa. **A.** Within T cell subset, expression of each molecule across mild, moderate, or severely inflamed oral mucosa following 6h PMA/I stimulation. n = 9 mild, 13 moderate, 11 severely inflamed oral mucosa tissues, unless noted based on minimum threshold for analysis of 25 cells. For Tregs, n = 8 mild; n = 11 moderate. No statistical differences across inflammation status for each marker. **B.** Expression of AREG, IFNγ, and IL-2 within each T cell subset; no statical differences across inflammation. **C.** Expression of each molecule within MAIT and γδ T cells. **D.** Representative gating of AREG+ GzmB+ populations within mildly inflamed oral mucosa (left) and co-expression of AREG+ GzmB+ among CD103+ CD8+ resident T cells (right). For MAIT cells, n = 7 mild, n = 8 moderate, n = 10 severe; γδ T cells, n = 7 mild, n = 10 severe. No significant differences by ordinary one-way ANOVA with Tukey’s multiple comparison test.

Similarly, the functional capacity of MAIT and γδ T cells was stable across inflammation states (***Figure 3C***). The mean, median, and data range are reported in Supplemental Table 2. Within several T cell subsets, we found co-expression of GzmB and AREG upon stimulation, which may indicate dual functional roles (cytotoxicity and tissue repair) can occur simultaneously within the same T cell. The highest frequency of GzmB and AREG co-expression was observed in CD8 CD103+ cells (***Figure 3D***), though these were also detected to a lesser extent in CD4 Tconv, CD4 Tregs, and innate-like subsets (∼1% within CD4 Tconv, Treg and MAIT cells, ∼4% in γδ T cells). Overall, these data suggest T cell subsets retained their functional capacity even in severely inflamed gingiva.

### T cells from lung and decidual-placental interference (DPI) differ based on tissue origin regardless of inflammation status

We next wanted to assess whether the phenotypic and functional stability of T cells across inflammation states is a gingiva-specific phenomenon or could also be extended to other mucosal tissues in healthy and inflamed states. We decided to examine the lung and the decidual-placental interface (DPI) as a set of mucosal tissues that are inherently very distinct from the oral mucosa in function and commensal exposure^42–44^. We assessed lung samples from deceased adult organ donors (healthy) along with resected lung tissue from patients with interstitial lung disease (ILD). Briefly, ILD is a group of pulmonary disorders causing inflammation and lung tissue scarring^45^. Tissue sampling was performed on both upper and lower lung lobes, which yielded nearly congruent profiles (*Supplemental Figure 6*); the upper lobe data are shown for the remainder of the figures. Decidual-placental interface (DPI) samples were taken from a cohort of term (37-41 weeks gestation), uncomplicated pregnancies (healthy) and a cohort of term pregnancies diagnosed with intra-amniotic inflammation (IAI). IAI, often referred to as chorioamnionitis, is typically an acute inflammatory response to a bacterial infection^46^.

First, we examined the composition of the T cell compartment in lung and DPI (***Figure 4A***). T cells isolated from DPI tissues displayed a higher frequency of naïve cells among all CD3+ CD45+, which are likely derived from the highly vascularized placental tissue. These naïve T cells were again excluded from manual gating analysis to facilitate a comparison with the memory phenotype T cells in other tissues. We noted a decrease in MAIT cell (p = 0.020) and γδ T cell (p = 0.133) abundance in inflamed over healthy lung, but did not observe significant inflammation-associated changes in the DPI (***Figure 4A***).

**Figure 4:**
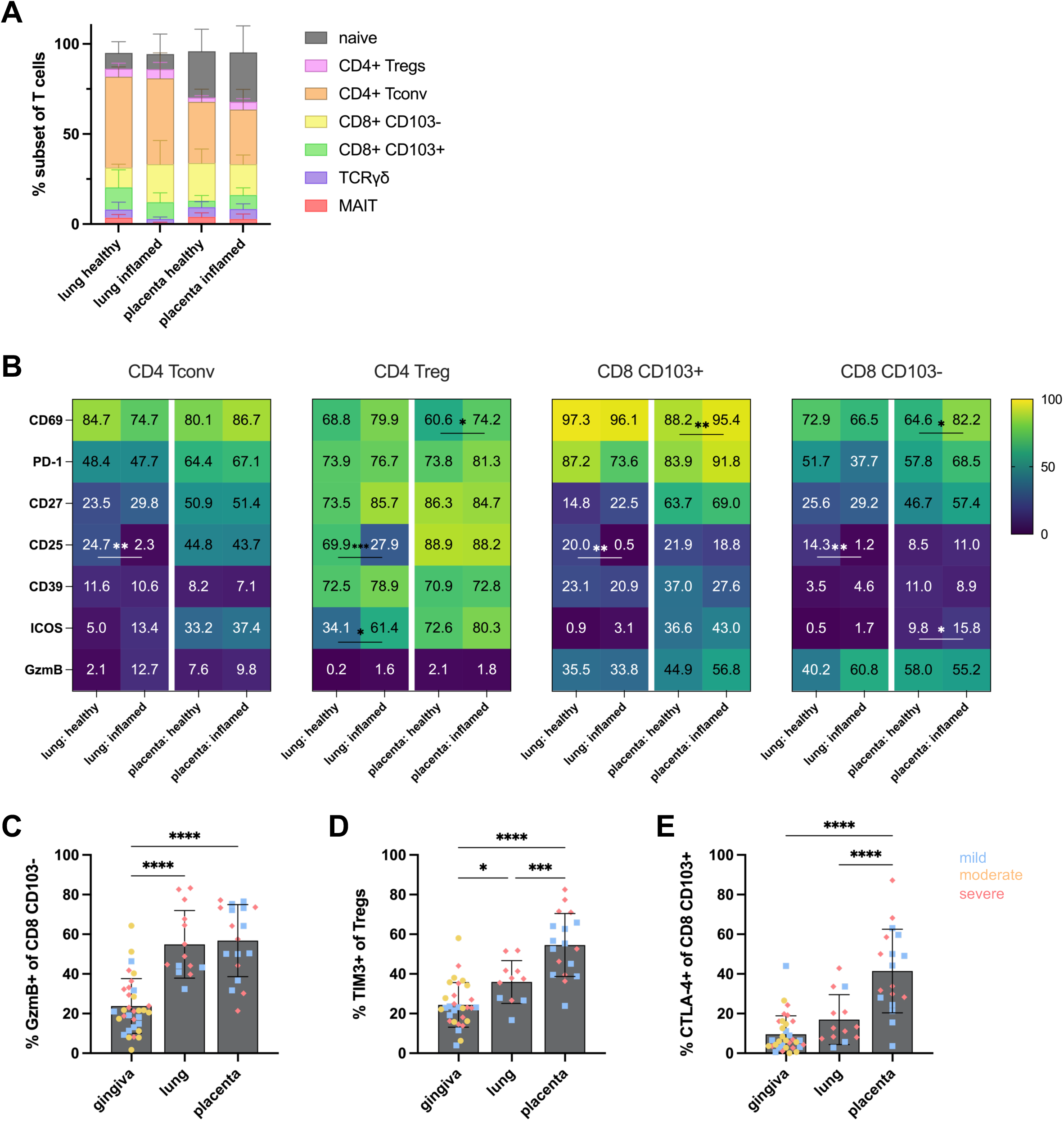
T cells from lung and decidual-placental interference (DPI) differ based on tissue origin regardless of inflammation status. **A.** Among all CD3+ CD45+ cells, frequency of indicated subset within each tissue and disease state. **B.** For conventional subsets (right to left; Tconv, Treg, CD103+ CD8+ and CD103- CD8+), frequency of activation markers by tissue and disease status. Unless noted, n = 4 healthy lung, n = 10 inflamed lung, n = 10 healthy placenta, n = 8 inflamed placenta. Some samples did not meet minimum threshold of 25 cells; for Tregs, n = 3 healthy lung; within CD103+, n = 3 healthy lung for all figure panels. Within one experiment, n = 8 inflamed lung samples for Tconv and Tregs for this and remaining panels. Statistical testing between tissues is excluded from heatmaps. **C.** Frequency of GzmB+ cells among CD8 CD103- across tissue and inflammation state (healthy/mild = blue squares, moderate gingiva = yellow circles, severe/inflamed = red diamonds). **D.** Frequency of TIM3+ Tregs. **E.** Frequency of CTLA-4+ cells among resident CD8 CD103+ cells. Pre-gated as indicated in Supplemental Figure 1 prior to subsetting and single-positive gating. *p < 0.05, **p < 0.01, ***p < 0.001, ****p < 0.0001 by ordinary one-way ANOVA with Tukey’s multiple comparison test.

Next, for each protein, the mean expression frequency is shown on all heatmaps in **Figure 4B**, Supplemental Figure 7, and the median and data range for each protein are shown in Supplemental Table 2. We noted an inflammation-associated, statistically significant increase in CD69+ T cells within CD4 Tregs, CD8 CD103+, and CD8 CD103- T cells in the placenta, but not the lung (***Figure 4B****, Supplemental Figure 7, statistical analysis across tissues excluded from heatmaps*). False discovery rate corrections for multiple comparisons of lung and placenta phenotype were also performed (*Supplemental Table 3*). A profound inflammation-associated phenotypic change occurred within T cells in the lung. We observed a dramatic reduction in CD25 expression on all T cell subsets in inflamed lungs (***Figure 4B***). Similarly to the oral mucosa, we did not observe an increase in GzmB in healthy vs. inflamed tissues (lung and DPI) (***Figure 4B***), and did not observe evidence for TCR-mediated activation, proliferation, or exhaustion (*Supplemental Figure 8*). We next wanted to further quantify how tissue origin (lung and DPI) vs. inflammation state affect protein expression of T cells. To accomplish this, we took the source of variation from 2-way ANOVA statistical testing (row factor and column factor p values with p < 0.05). We observed that tissue origin had a greater impact on subset composition and phenotypic differences than inflammation status. (*Supplementary Table 2, markers displayed on heatmaps in Supplemental Figure 7*).

So far, the data indicate that tissue origin predominantly shaped the general phenotypic profile of T cell subsets. We next combined all 3 tissue types for our subsequent analysis steps. To further visualize this relationship of phenotype and tissue location vs. inflammation state across all three tissue sets, we grouped all datapoints from each tissue (irrespective of inflammation state), color-coded each datapoint according to inflammation state, and compared the expression of T cell surface markers across gingiva, lung, and placenta within conventional T cell subsets (***Figures 4C-E***). As highlighted in previous figures, we did not observe inflammation-associated changes in GzmB expression in CD8 T cell subsets, however there was a significant difference in GzmB expression in CD8 CD103- T cells in the gingiva compared to lung and DPI (***Figure 4C***). Among these tissue sets, the gingiva is presumably the tissue that is most consistently and substantially exposed to microbes and antigen yet still displayed the lowest baseline GzmB expression in CD8 CD103- T cells. Similarly, the frequency of TIM3+ Tregs was lowest in the gingiva (***Figure 4D***). TIM3 expression by Tregs is typically observed in highly activated Tregs in the tumor microenvironment^47^. In our dataset, we observed a nearly 3x increase in frequency of TIM3 expression in Tregs from the DPI compared to gingiva (***Figure 4D***). Together, these data suggest that the DPI, although typically considered to be a sterile tissue environment^42^, shows signs of being concomitantly activated and suppressive. In line with this notion, we also observed the highest frequency of CTLA4+ expressing CD8 CD103+ T cells in the DPI (***Figure 4E***). Together, these data highlight that tissue-specific phenotypic adaptations in the T cell compartment across these mucosal tissues were maintained during states of inflammation. We next wanted to test if the functional capacity of these T cell subsets was similarly maintained in all tissues during inflammation.

### Functional capacity of T cells is primarily shaped by tissue location

As a first step, we examined the functional capacity of T cell subsets across lung and DPI during health and disease. We found that the expression pattern of cytokines was relatively stable across inflammatory states. Two statistically significant changes between healthy and inflamed lung were noted in CD4 Tconv; a decrease in IL-17A and IL-17F expressing cells in inflamed lungs (***Figure 5A****, Supplemental Figure 9, statistical analysis across tissues excluded from heatmaps*). Full statistical testing (FDR corrections for multiple comparisons) is detailed in Supplementary Table 3.

**Figure 5:**
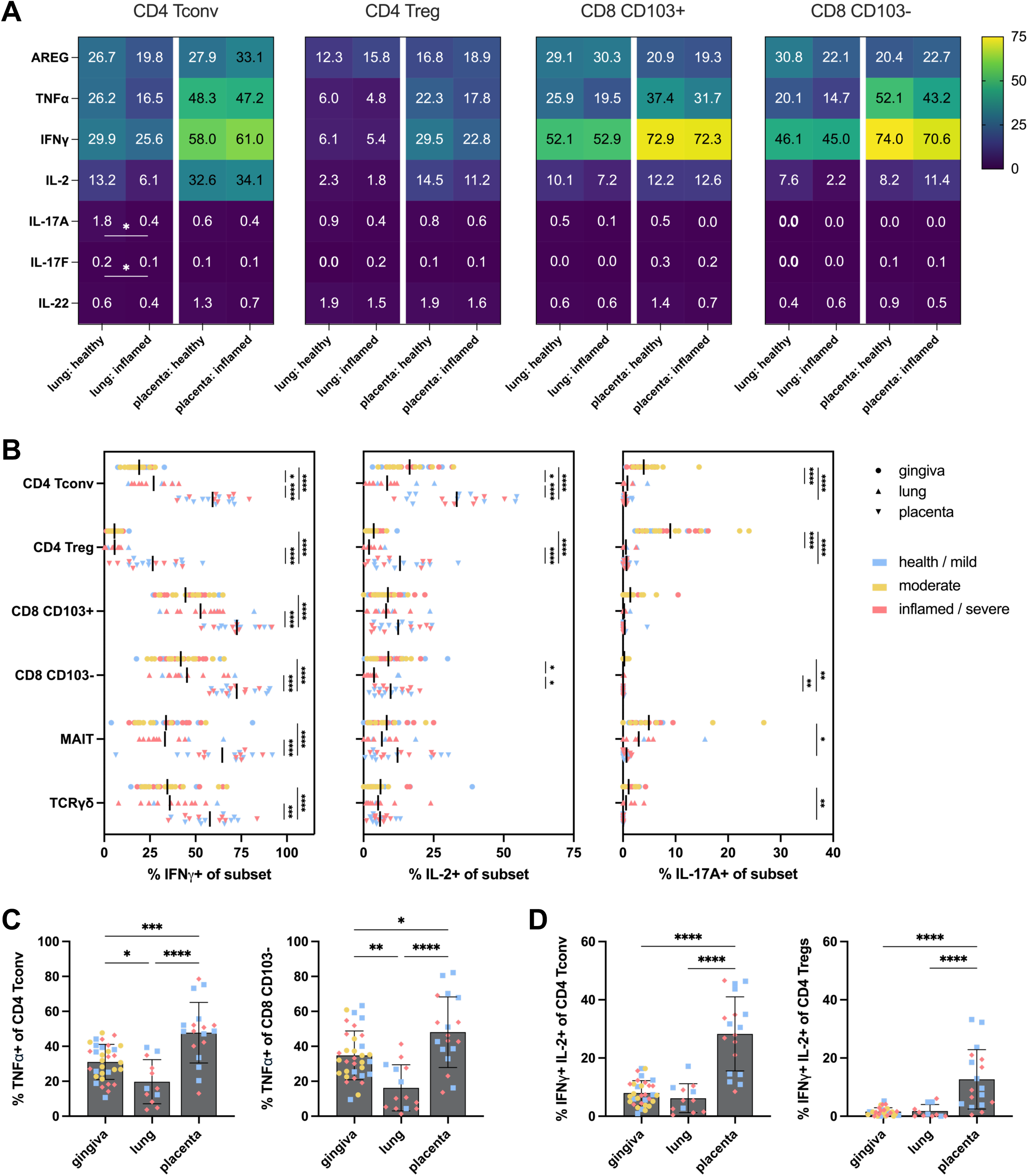
Functional capacity of T cells is primarily shaped by tissue location. **A.** Upon 6hr stimulation with PMA/ionomycin, expression of cytokines and tissue healing markers across T cell subsets in the lung and placenta. Unless noted, n = 4 healthy lung, n = 10 inflamed lung, n = 10 healthy placenta, n = 8 inflamed placenta. For Tconv and Tregs, n = 8 inflamed lung samples for this and subsequent figure panels. Statistical testing between tissues excluded from heatmaps. **B.** Within each T cell subset, expression of IFNγ (left), IL-2 (middle), and IL-17A (right). For all panels, circles = gingiva, upright triangle = lung, upside down triangle = placenta; for this and subsequent figure panels, healthy/mild = blue, moderate gingiva = yellow, severe/inflamed = red. For MAIT cells, n = 3 healthy lung and n = 9 inflamed lung samples. **C.** TNFα expression across tissues in CD4 Tconv (left) and CD8 CD103- (right), pre-gated based on Supplemental Figure 1 scheme. **D.** Frequencies of IFNγ+ IL-2+ across tissues in CD4 Tconv (left) and Tregs (right). *p < 0.05, **p < 0.01, ***p < 0.001, ****p < 0.0001 by ordinary one-way ANOVA with Tukey’s multiple comparison test.

We next compared the functional capacity across all three tissue sites and inflammation status, (***Figure 5B****, gingiva – top, circles; lung – middle, triangles; placenta – bottom, diamonds*). These data highlight that the greatest capacity for T cell IFNγ production (***Figure 5B***, left panel) and IL-2 (***Figure 5B***, middle panel) was observed in the DPI. TNFα was also produced in the highest quantities by T cells from the DPI and lowest in the lung (***Figure 5C***). In contrast, IL-17A expression was essentially absent in DPI T cell subsets and most prominent in the oral mucosa (***Figure 5B***, right panel). Differences in the functional capacity of CD4 Tregs particularly stood out (***Figure 5B***). The oral mucosa contained a large fraction of Tregs that expressed IL-17A following stimulation (***Figure 5B***, right panel), while Tregs from the placenta expressed IFNγ (***Figure 5B***, left panel). Of note, we observed co-expression of IFNγ and IL-2 for both CD4 Tconv and Tregs from the DPI (***Figure 5D***) thus potentially suggesting that some FoxP3+ CD127- T cells may be activated CD4 Tconv (explored further in the discussion). Finally, it is noteworthy that the bias towards certain functional properties in each tissue set was mirrored across several T cell subsets. For example, gingival Tregs were the most potent IL-17 producers, and this was true for CD4 T cell subsets, but also MAIT cells and γδ T cells (***Figure 5B***) regardless of inflammation status. Overall, these data support the notion that the functional capacity of T cell subsets is linked to tissue location, and that this can be maintained during inflammatory disease.

### Functional capacity of T cells across tissues and inflammation state by dimensionality reduction analysis

Our analysis approach so far was focused on specifically examining select biomarkers of activation and functional properties. We next wanted to compare tissues in a more unsupervised analysis approach. Of note, we performed this analysis separately for stimulated and unstimulated samples. Stimulation – as expected – altered the expression of some cell surface biomarkers. We reasoned that the presence vs. absence of stimulation-induced changes could also be informative and included all phenotypic and functional markers in the antibody staining panel in this PCA analysis (TIM3, CD39, ICOS, CD25, CD28, CD27, PD1, Ki67, IL1R1, CD137, CD40L, CTLA4, IL-2, IL-22, AREG, IL-17A, IFNγ, GzmB, TNFα, IL-17F). We first used principal component analysis (PCA) to compare the effect of tissue origin vs. inflammation status on T cell functional capacity (***Figures 6A-D****, Supplemental Figure 10)*. Tissue-based differences in functional capacity were particularly apparent in CD4 Tconv and CD8 CD103+ T cells (***Figures 6A-B***) and were somewhat less pronounced in MAIT and γδ T cells (***Figures 6C-D***). In contrast, for all subsets, there was near congruent clustering based on inflammation status.

**Figure 6:**
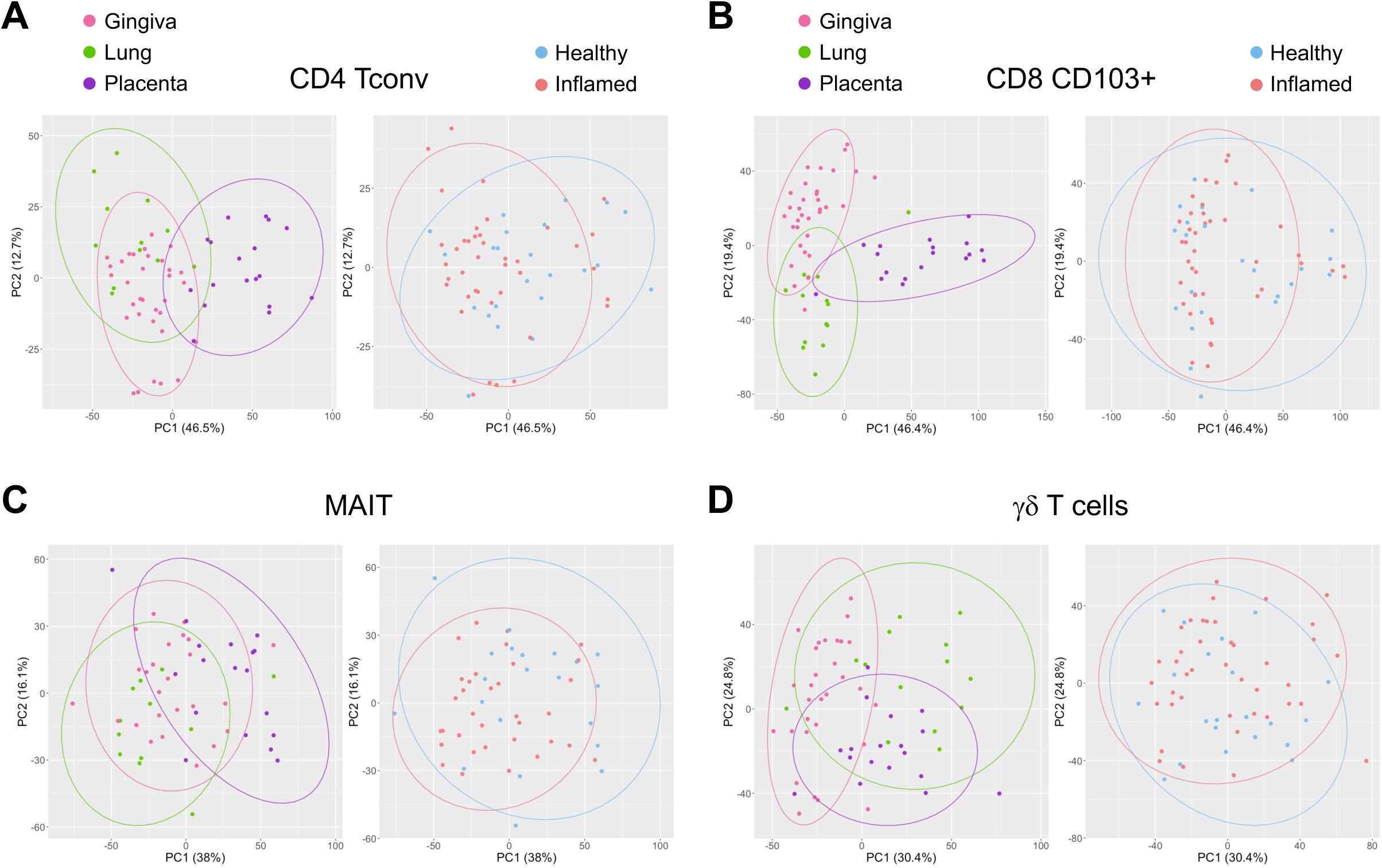
Functional capacity of T cells across tissues and inflammation state by dimensionality reduction analysis. Principal components determined by frequency of positive cells within gates from the following parameters: TIM3, CD39, ICOS, CD25, CD28, CD27, PD1, Ki67, IL1R1, CD137, CD40L, CTLA4, IL-2, IL-22, AREG, IL-17A, IFNγ, GzmB, TNFα, IL-17F. PCA plots of flow cytometry data following 6-hour PMA/Ionomycin stimulation from gingiva (pink), lung (green), and placenta (purple) during health (blue) and inflammation (red) organized by tissue type (left) and inflammation status (right) of **A.** CD4 Tconv, **B.** CD8 CD103+, **C.** MAIT cells, **D.** γδ T cells. Moderately inflamed oral mucosa is grouped with inflammation category. Ellipses represent 95% confidence interval.

When determining which parameters are contributing most to determining the principal components (loading scores), upon stimulation, changes in IFNγ expression contributed the greatest for all T cells. Differences in expression of the co-stimulatory molecules CD27 and CD28 were also significant contributors to the PCA. Importantly, similar findings were observed when PCA was performed with unstimulated samples directly *ex vivo* (*Supplemental Figure 11)*. Taken together, an unsupervised analysis approach further underlines the notion that tissue inflammation is not sufficient to disrupt tissue-specific adaptation of the T cell compartment.

### An unsupervised analysis approach to compare the T cell compartment across tissues and inflammation

We next wanted to use an unsupervised approach to assess the relative abundance of T cells with functional similarity across tissue and inflammation state. An unsupervised approach also allows for discovery of phenotypically/functionally unique populations potentially missed by manual gating of each marker using single- or double-positive gates. Of note, for this approach, we included naïve T cells in the analysis as a reference population. Briefly, T cell subsets were first defined by manual gating. We then performed scaling/normalization and defined global clusters across all samples (described further in methods). We assessed each T cell subset using this approach with tissues analyzed directly *ex vivo* and following stimulation. Given the profound differences of Treg functional capacity between tissues, we first focused our analysis on stimulated Tregs. We examined cluster frequencies across tissues (gingiva, pink; lung, green; placenta, purple) (***Figure 7A***, *top row panels*) and across inflammation status (healthy, blue; inflamed, red) (***Figure 7A***, *bottom left panels*).

**Figure 7:**
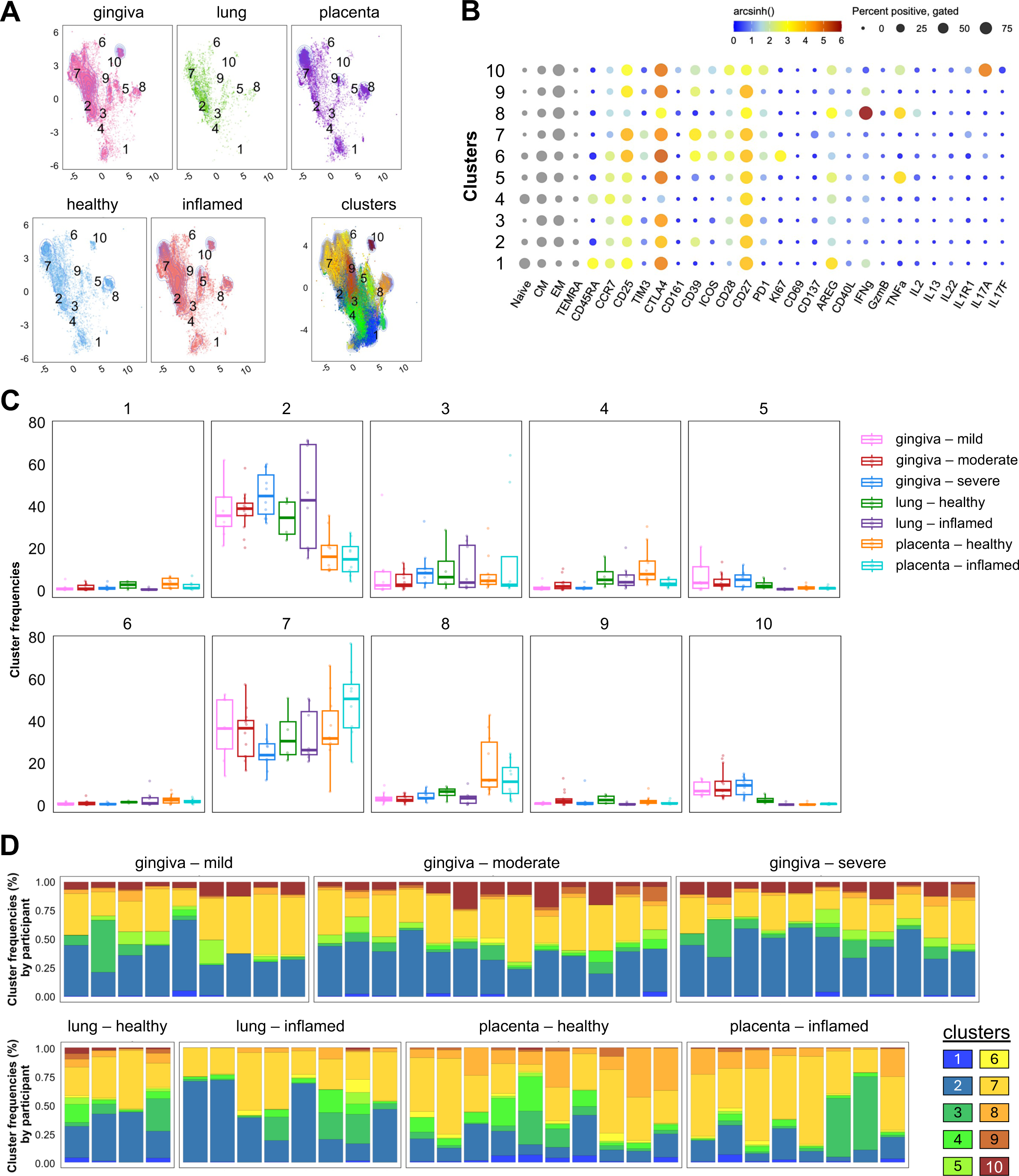
An unsupervised analysis approach to compare the T cell compartment across tissues and inflammation. The following markers were used to develop clusters: CD161, CD39, CCR7, ICOS, CD25, IL-17F, CD28, CD27, CD45RA, PD-1, Ki67, CD69, TIM3, IFNγ, CTLA4, IL-13, IL-22, GzmB, TNFα, IL1R1, CD137, AREG, CD40L, IL-17A, CD45RO, IL-2. **A.** UMAPs of clusters organized by tissue origin and inflammation status. **B.** Bubble plots of expression of each marker within each cluster among Tregs upon stimulation. Resolution = 0.75. **C.** Box plots of frequency of each cluster relative to CD4 Treg counts out of total T cells from each sample. **D.** Stacked bar graphs representing cluster breakdown for individual samples.

We next assessed the characteristics of the observed ten distinct population clusters using a bubble plot (***Figure 7B***). The bubble plot highlighted that clusters 1 and 4 were comprised of naïve cells based on the expression of CD45RA+ CCR7+, whereas the remaining clusters consisted of memory subsets. The relative abundance of each cluster combined for all donors is displayed in **Figure 7C**, and then further broken down by single donor in **Figure 7D**. Tregs that expressed the cytokines IFNγ and IL-17A were in clusters 8 and 10, and – in line with our observations from manual gating – most abundant in placenta and oral mucosa, respectively. Overall, these data indicate that our manual gating analysis captured differences in Treg functional capacity across tissues (***Figure 5B****, Supplemental Figure 9*). We next performed this analysis for other T cell subsets. Clustering for CD4 Tconv cells across tissue and inflammation is shown in *Supplemental Figure 12*; the results are also in line with our manual gating analysis. To summarize all clustering data for other T cell subsets *ex vivo* and upon stimulation, we used clustering frequencies to generate PCA plots (*Supplemental Figure 13-14*). When cluster frequencies were used to determine principal components, distinct tissues (gingiva, lung, and DPI) form unique groups, whereas healthy and inflamed tissues overlap. Overall, our unsupervised analysis approach further highlights that clusters show tissue-specific characteristics with minimal alterations due to inflammation status.

In summary, across all analysis methods, we found that tissue-specific adaptations of T cell subsets were maintained during tissue inflammation.

## Discussion

Our main goal was to assess the extent to which tissue-specific adaptation of the T cell compartment is altered during tissue inflammation. We initially sought to address this question by examining oral mucosal (gingiva) tissue, but subsequently extended our study to include other mucosal tissues to ensure that our conclusions would not be limited to a single tissue. In the gingiva, we observed that markers indicative of inflammation-driven T cell activation did not significantly change in most T cell populations, including MAIT and γδ T cells, as inflammation severity of the gingiva increased. We considered that this could be due to the baseline level of inflammation that is present even in healthy gingival tissues^35^, which could yield unique regulatory mechanisms in the tissue. Since the oral mucosa is colonized by a dynamic microbiome and continuously exposed to dietary and microbial antigens^35^, we also examined the expression of biomarker pairs that indicate TCR-mediated activation. ICOS and PD-1, and CD39 and PD-1 have been shown to be indicative of recent TCR activation for human CD4 and CD8 T cells, respectively^39–41^. Neither of these biomarker pairs indicated an association of a TCR-mediated T cell response in CD4 Tconv or a CD8 T cell subset gingiva with increased severity in inflammation. Similarly, Ki67 expression, an indicator of cell proliferation, and expression of PD-1 and TIM3, indictors of T cell exhaustion, did not change with increased inflammation in the gingiva. Overall, our observations highlight that the overall composition and the phenotypic properties of T cell subsets in the gingiva were largely maintained at various states of inflammation.

Because of the unique physiological demands of the gingiva, including but not limited to mastication, we asked if the T cell compartment would be similarly resistant to inflammation-mediated changes in other mucosal tissues, specifically the lung and the decidual-placental interface (DPI). CD4 Tconv and CD8 T cell subsets in the healthy lung did not express biomarker pairs of TCR activation. However, in the DPI we found that a substantial fraction of CD4 and CD8 T cell subsets co-expressed ICOS and PD-1, and CD39 and PD-1, respectively (*Supplemental Figure 8*). These data could suggest that T cells in the DPI encounter their antigen more frequently than T cell in the gingiva and lung. This could potentially be related to the presence of fetal-derived, semi-allogenic tissue (the placenta is made of fetal-derived trophoblasts) or could indicate that recently activated T cells migrate to the DPI to provide protection. An in-depth analysis of the TCR repertoire in the DPI compared to other tissues may provide a more definitive answer in future studies. Finally, ICOS, CD39 and PD-1 are all expressed by highly activated Tregs, and expression of these markers was found on a large fraction of Tregs across all tissues and inflammation states.

Given the presence of highly activated Tregs in all tissues, we explored whether Tregs displayed functional differences. We found that in gingival tissues, regardless of inflammation status, about 10% of Tregs expressed IL-17A. IL-17+ Treg have been previously described and are thought to maintain their ability to suppress other T cells, while secreting IL-17 in response to TCR-mediated activation, or IL-6 and IL-1β stimulation^48,49^. Importantly, both IL-6 and IL-1β are expressed in healthy human gingiva^50,51^. In contrast, we did not observe IL-17+ Tregs in lung or DPI. However, we found that a fraction of Tregs in both healthy and inflamed DPI could express IFNγ, while none of the Tregs in other tissues expressed IFNγ following stimulation. These cells could be Th1-like Tregs that have so far been mainly described in peripheral blood in context of human autoimmune diseases and are thought to be hypo-suppressive^52^. Thus, our observation could indicate that these Tregs also exist in a healthy human tissue (albeit from a unique environment that contains a semi-allogeneic tissue), but it important to consider that these could potentially be recently activated Th1 cells that temporarily express FoxP3^53–55^. Alternatively, these IFN γ^+^ Tregs could be specialized to suppress Th1 responses as previously shown in a mouse model system^56^. Follow up studies will be required to distinguish between these possibilities and transcriptionally define this DPI-enriched Treg population. Overall, tissue-associated phenotypes and functional properties were evident for Tregs, while inflammation-associated changes were limited in our cross-tissue and inflammation-state comparison.

One of the most prominent phenotypic changes that we observed between a healthy and inflamed tissue was the striking decrease in CD25 expression on T cell subsets in the inflamed lung (Figure 4C, *Supplemental Figure 7*). Our PBMC batch control (*Supplemental Figure 3*) allowed us to rule out technical reasons for these phenotype (wrong titer of the antibody, etc.). Previous reports indicated an increase in serum soluble CD25 levels in individuals with ILD (both granulomatous-lymphocytic and rheumatoid-arthritis associated ILD)^57,58^. ILD has also been clinically associated with increased matrix metalloproteinase (MMP) expression in serum^59,60^, which cleaves CD25 from the surface of T cells. If there is a local increase in MMP activity in ILD lungs, then this could provide an explanation of the decreased CD25 during ILD in all T cell subsets.

We specifically looked for functional changes that have previously been associated with inflammation status. Studies using gingival tissue indicated a potential association between inflammation and the abundance of Th17 cells^61,62^. We found that the overall composition of the T cell compartment, including the frequency of CD4 T cells that make IL-17A or IL-17F following stimulation, is relatively stable even during inflammation. It is important to keep in mind that a hallmark feature of inflammation is the recruitment of immune cell populations to a site of inflammation, and that inflammation of the gingiva is associated with an increase in T cell infiltration^63^. Thus, while the overall number of T cells with a specific function increases in inflamed tissues, this is not necessarily associated with a skewing in functional properties within the T cell compartment. Overall, the composition of the T cell compartment was consistent between mildly to severely inflamed gingiva, with few subsets showing an increase or decrease with inflammation. Specifically, we observed a reduction in resident CD8 CD103+ T cells within more severely inflamed gingival tissue with a concomitant proportional increase in circulating CD8 CD103- T cells. Of note, our study does not suggest that stability is always maintained. This stable maintenance of function would presumably change, at least temporarily, in disease-specific conditions with an associated antigen-specific T cell response, such as the Th17-driven response against *Candida albicans* and *Staphylococcus aureus*^64^.

Overall, a state of tissue inflammation does not per se override tissue-specific features of the T cell compartment. Importantly, our observations for the T cell compartment do not necessarily extend to inflammation-associated alterations of other cell types, particularly innate immune cells, in these tissues. In the gingiva, it is well documented that neutrophil infiltration is associated with acute inflammation^61,65,66^. In the lungs, similar innate subsets of macrophages and neutrophils have been reported to drive inflammatory responses in mice and humans^67,68^, and acute chorioamnionitis has also been defined by neutrophil infiltration^69^. Again, it is important to keep in mind that specific diseases can profoundly impact T cell phenotype and function during inflammation. Thus, while our data indicate that tissue inflammation is not inherently sufficient to override tissue-specific T cell adaptation, different disease settings may display profound T cell changes such as previously observed in the context of Hepatitis A infection, which leads to increased IL-15 availability and bystander activation of CD8 T cells^27^.

Finally, we want to highlight that we used a brief PMA/Ionomycin stimulation condition to examine the functional capacity of T cells. We reasoned that the broadly activating nature of this stimulus will reveal which proteins T cell subsets are able to express, while keeping in mind that it does not reveal which proteins are secreted in situ. This approach indicated that T cell subsets retained their ability to express AREG even in chronically inflamed tissue such as the ILD lung and severely inflamed gingiva. Whether AREG is actually secreted *in situ* by T cells to promote epithelial cell recovery in the inflamed gingiva is unclear. CD8 T cells appear to require a TCR signal to secret AREG^34^ and given the lack of evidence for CD8 T cell activation (including TCR-mediated T cell activation) in these tissues, *in situ* AREG secretion by CD8 T cells seems unlikely. However, given that the functional capacity is maintained in inflamed tissues, novel therapies could target AREG to elicit expression or other specific functional responses.

In summary, we report that T cells are able to maintain their tissue-specific core identities during states of tissue inflammation. By gaining a more complete understanding of how inflammation affects the tissue-specific T cell landscape, therapeutic approaches can be designed to develop treatments to restore homeostasis with tissue-specific requirements in mind.

## Material and Methods

### Approvals & Study Population

#### Tissue Samples and Protocols

All protocols used in this study were approved by the Institutional Review Board (IRB #0008335, 00008263, 00004036, 00001636). Signed informed consent was obtained from all participants. A list of all samples, along with relevant clinical information can be found in Supplementary Table 1. Each sample is from a unique donor.

#### Oral Mucosa

Oral mucosa samples were obtained from surgically discarded tissue from routine surgeries, such as osseous surgery, functional or esthetic crown lengthening procedures, tooth extractions, or impacted tooth uncover procedures. Mild inflammation (clinical health, mild to moderate gingivitis), moderate inflammation (severe gingivitis, stage I, and stage II periodontitis), and severe inflammation (stage III and IV periodontitis).

#### Lung

Healthy lung samples were redirected transplants from deceased organ donors. Donors without history of lung disease, infection, and mechanical ventilation for >7 days were eligible for screening. Inflamed lung samples (interstitial lung disease, ILD) were obtained from patients undergoing lung transplant. Explant lungs and hilar lymph nodes were obtained from patients with ILD undergoing lung transplant at the University of Washington.

#### Decidual-Placenta Interface

Both healthy and acutely inflamed (chorioamnionitis) third trimester placentas were obtained following vaginal delivery or C-section, following labor or no labor (detailed in Supplementary Table 1).

#### Technical Control Sample

A consistent leukopak peripheral blood mononuclear cell (PBMC) sample was used throughout each flow cytometry batch and assessed alongside experimental samples to ensure longitudinal control. Leukopak (4510-01 Ful Leuko Pack) obtained through Bloodworks Bio (Seattle, WA).

### Isolation of Lymphocytes from Tissues

#### Oral Mucosa

Following procedures, mucosal tissue was priority shipped overnight with cold packs to Fred Hutch Cancer Center (FHCC). Upon arrival, tissue was washed with complete media (RPMI1640 (11875-093, Gibco) with penicillin, streptomycin (15140-122, Gibco), and 10% fetal bovine serum (FBS, PS-FB2, Peak Serum)). An optimized protocol was adapted from reference^70^, originally developed for processing solid tumors. Tissue was placed in a petri dish and minced with a scalpel, then enzymatically digested (Collagenase II (C6885, Sigma-Aldrich), DNAse (20 units/ml, #D502, Sigm) in complete media with reduced 7.5% FBS (RP7.5) in a 25 mL volume at 37°C for 40 minutes while shaking. Following digest, the sample was passed through a needle (16×1 ½ Blunt, 302833, BD) 10 times prior to passing through a 100-micron cell strainer. Remaining tissue pieces were mashed using a syringe plunger, washed with complete media, and frozen in 1 mL Recovery Cell Culture Freezing Medium (12648010, Gibco) and stored in liquid nitrogen until future experiments. Cell viability and abundance were quantified prior to freezing by staining a small sample aliquot (Guava antibodies 7-AAD, CD19, and CD45) for 15 minutes, resuspension in 1% PFA, and collection on the Guava easyCyte. Oral mucosa samples in this study contained between 731-52,195 T cells, as analyzed by flow cytometry.

#### Lung

Lung tissue was processed in a similar fashion to the gingiva using the RP7.5 digestion for 50 minutes at 37°C while shaking at 225 RPM. Subsequently cells were washed with RP10, strained through a 70µm filter and enumerated. Lung tissue samples contained between 1,537-62,719 T cells.

#### Decidual-Placenta Interface

Tissue was processed as previously described^71^. Briefly, intact placentas were biopsied up to a 0.5 cm depth on the maternal facing surface of the placenta including both decidua basalis and placenta villi. Tissue was digested in digestion media (above) and incubated at 37°C for 60 minutes with agitation. Concentration for digest was RP7.5 + 700 U/mL Collagenase Type II (Sigma-Aldrich) and 200 U/mL DNase. Cells were filtered and washed with media. Red blood cells were removed using ACK lysis buffer, followed by resuspension with media. Maternal blood was collected in tubes containing the anti-coagulant acid citric dextrose. DPI tissue samples contained between 10,303-102,650 T cells.

#### Technical Control Sample

Leukopak was processed according to StemCell Technologies protocol (https://www.stemcell.com/leukopak-processing-protocol.html). Two blood cell lysis steps at room temperature and two platelet washes were performed prior to counting and freezing 20 million cells per aliquot.

### Stimulation Assay

As briefly outlined in Figure 1A, upon experimentation, cryo-preserved single-cell suspension isolated from tissues or blood (outlined above) were thawed in a 37°C water bath. 12,000 to 2 million cells per condition were plated in a U- or V-bottom 96 well plate in 200 uL complete media. For assessing phenotype directly *ex vivo*, cells were left untreated for 2 hours at 37°C. GolgiStop (Monensin, 554724, BD) and GolgiPlug (Protein Transport Inhibitor (Containing Brefeldin A), 555029, BD) were then added per manufacturer recommendations to compare directly to stimulated samples and was returned to the 37°C incubator for 4 more hours (total of 6 hours). During functional stimulation assays, samples were similarly thawed and plated. 200 uL complete media supplemented with PMA (100 ng/mL, P8139, Sigma) and ionomycin (1000 ng/mL, I9657, Sigma) for 6 hours, including the addition of Golgi stop/plug 2 hours after beginning the stimulation. Upon completion, samples were stained for analysis by flow cytometry.

### Spectral Flow Cytometry

#### Staining

For all experiments, upon completion of the assay, samples were stained previously described^72^. Prior to experimentation, the spectral flow panel was tittered, optimized, and tested to ensure proper design and maximal information could be obtained from each sample, as detailed in reference^73^. All staining steps were performed in the dark at room temperature in a 96-bottom plate. Cells were incubated with FcX block (422302, Biolegend) and Zombie NIR dye (422302, Biolegend) in PBS (14190250, Gibco) for 15 minutes. Antibody master mixes were spun down at 10,000xg for 4 minutes prior to staining to remove aggregates. Following live/dead staining, cells were washed, following by surface staining in 50uL (antibody mix in PBS + 2% FBS and 10% brilliant stain buffer (563794, BD)) for 20 minutes. Cells were then washed twice prior to fix/perm (100 uL, 554722, BD) according to manufacturer guidelines for 30 minutes. After fixation, cells were stained intracellularly for select markers noted in the panel (Supplementary Figure 1) for 30 minutes, assembled in perm/wash buffe and 10% BSB. Finally, cells were washed twice with perm/wash and resuspended in PBS + 2% FBS and stored at 4°C until acquisition the following day on the BD Discover S8.

#### Controls

Single-stain controls were optimized for individual markers by staining either unstimulated PBMCs, PBMCs following a 6-hour PMA/ionomycin stimulation, or unstimulated control tissue samples (generally tonsil tumor infiltrate or regional lymph nodes). For two markers expressed in low abundance in PBMC and tissue (IL-13 and IL-22), capture beads were used for single-stain control. Control samples were taken through an identical staining protocol to experimental samples, described above. A reference sample was also used within each experiment as a longitudinal control (*Supplemental Figure 3*).

#### Cytometer

Data from all experiments was collected with the BD Discover S8 and the FACSChorus software platform (BD Biosciences). The S8 is equipped with 5 lasers and 78 detectors. Compensation controls were prepared and used for each individual experiment, aside from one run containing ILD 11 and 12. Upon collection, data was exported (FCS 3.1 format) and analyzed using FlowJo version 10.10.0. Files were analyzed in batches by collection day, with minor gating changes across different batches.

### Unsupervised Clustering

Upon acquisition on the Discover S8 (BD), samples were gated (*Supplemental Figure 1*). Following gating of time, lymphocytes, single cells, and live cells, T cells were gated (CD3+ CD45+) and exported as a new .fcs file in FlowJo. Files were also stripped of parameters out of scope for this analysis, focusing on time, size channels, and those containing a fluorochrome (*listed in Supplementary Figure 1*). Upon creation of files containing only T cells, separate streams were created for each relevant T cell subset assessed in this study: γδ T cells (TCRγδ+), MAIT cells (CD161+ Vα7.2+), all CD4 (CD8-), Tconv (FoxP3-), Tregs (FoxP3+ CD127-), all CD8 (CD8+ CD4-), resident CD8 cells (CD103+), and circulating CD8 cells (CD103-). Naïve cells were included in clustering analysis for all tissues.

We included additional samples in our clustering analysis as reference tissues for tissue-enriched T cell subsets (*Supplemental Table 1, gray lines*). Lung and associated lymph nodes, along with maternal blood from placental donors were included in multiple batches. We also included tumor infiltrates from head and neck squamous cell carcinoma (HNSCC) and associated lymph nodes. One experimental batch did not include marker CD127 and was excluded from the Tconv and Treg clustering analyses (n = 1 pediatric lung and associated LNs, 2 inflamed lungs and associated LNs, 2 tumor infiltrate samples, and 8 tumor associated LNs).

For each clustering stream, we used up to 20,000 T cells per sample. No minimum cell threshold was utilized. Actual T cell counts for individual samples and primary clustering streams were considered for count analysis. The following markers were used to develop clusters: CD161, CD39, CCR7, ICOS, CD25, IL-17F, CD28, CD27, CD45RA, PD-1, Ki67, CD69, TIM3, IFNγ, CTLA4, IL-13, IL-22, GzmB, TNFα, IL1R1, CD137, AREG, CD40L, IL-17A, CD45RO, IL-2.

Lineage markers and QC parameters were excluded from clustering analysis: CD3, CD45, CD4, CD8, FoxP3, CD127 or CXCR5, CD103, TCRγδ, Vα7.2, Live/Dead, Autofluorescence. Spectre^74^ was used to determine appropriate arcsinh transformation values for each parameter.

Repeated clustering at the single cell level – necessary to identify and factor out technical noise, or perceived batch effect – is computationally intractable at the scale of this study. Therefore, within each of the streams, to arrive at globally defined clusters across all batches, we cluster the centers of clusters from batch-specific clusterings^75,76^. Specifically, we ‘over’-cluster within each batch by first clustering, then subclustering the cells within each cluster. Next, we compute the centroids of all (sub)clusters in each batch and cluster these (sub)cluster centroids (or “metacells’’ or “supercells”) to arrive at a global (meta)clustering assignment for each metacell from each batch^77^. At each stage we use the Rphenograph algorithm, with Leiden clustering of shared nearest-neighbor graphs^78,79^. To address moderate drift across batches in some parameters, we first align repeated control median transformed intensity values across batches for select parameters, and next apply the same batch- and parameter-specific shifts to the samples before metaclustering. We use Uniform Manifold Approximate Projection (UMAP) for dimension reduction and data visualization within an iterative process of adjusting for perceived batch effects, optimized by refining the metaclustering at a fixed Leiden resolution, before pushing resolution higher^80^.

### Statistical Analyses

Statistical testing was performed using GraphPad Prism Software version 10.5. Testing us noted within figure legends. All bar graphs represent the mean with standard deviation. Heatmaps display the mean for each marker for a given sample group. For oral mucosa samples (*Figures 1-3*), significance was determined by testing using ordinary one-way ANOVA with Tukey’s multiple comparisons test. For lung and placenta data (*Figures 4-5*), 2-way ANOVA was utilized with Tukey’s multiple comparisons if comparing two tissues across inflammation states, or one-way ANOVA if comparing with three tissues with grouped inflammation. All data using one-way ANOVA was also assessed using the Benjamini-Hochberg false discovery rate (FDR) correction with a rate of 10% and maintained statistical significance. Supplementary Table 2 used 2-way ANOVA to compare lung and placenta across inflammation; the Benjamini-Hochberg FDR correction with a rate of 10% was utilized.

## Resource Availability

Raw flow cytometry files (.fcs) were submitted to ImmPort and assigned as Study SDY3484.

## Acknowledgements

We thank the Flow Cytometry Shared Resources Core at Fred Hutch Cancer Center. We thank M. Quinn Peters who provided and assisted with the R script used for PCA and Veronica Davé who consulted on proper statical testing and corrections to be used for these analyses. We also thank all members of the Prlic Lab for critical feedback and discussion.

## Author contributions

Conceptualization: N.B.P., M.P. Methodology: N.B.P., H. R. M., M.P. Conducting experiments: N.B.P., E. D. V., A. K. T., C. S. D., G. D. Formal analysis: N.B.P., H. R. M., M.P. Writing; original draft: N.B.P., M.P. Writing; review and editing: all authors. Visualization: N.B.P., H. R. M., M.P. Funding acquisition: N.B.P, M.P. Supervision: M.P.

## Funding

NIH grants R56DE032009, R01AI123323 and R01AI179712 (to M.P) and funding from the Oral Mucosa Immunology Consortium at the University of Nebraska (to N.B.P, Windsweep Farm Scholar) supported this work.

## Declarations of interest

The authors declare no competing interests.

## Supplemental Information

Document S1. Figures S1-S14.

Tables S1-3 contain tissue information, data range of heatmaps, and statistical testing.

## Notes

### Competing Interest Statement

The authors have declared no competing interest.

### Summary of Updates

Updated Figure 7B - bubble plot missing in previous version

