## Supplemental Figures for "Tissue-specific adaptation of human T cells is preserved during tissue inflammation"

### Supplemental Figure 1

**A**

| Fluorochrome | Antigen | Clone | Vendor | Dilution | Staining |
|---|---|---|---|---|---|
| Spark UV 387 | CD4 | SK3 | Biolegend | 1: 80 | Intracellular |
| BUV395 | CD8a | RPA-T8 | BD | 1: 80 | Surface |
| UV Live Dead | Autofluorescence | NA | NA | NA | NA |
| BUV496 | CD3 | UCHT1 | BD | 1: 40 | Intracellular |
| BUV563 | CD161 | DX12 | BD | 1: 20 | Surface |
| BUV615 | CD39 | TU66 | BD | 1: 20 | Surface |
| BUV661 | CCR7 | 2-L1-A | BD | 1: 10 | Surface |
| BUV737 | ICOS | DX29 | BD | 1: 160 | Surface |
| BUV805 | CD45 | HI30 | BD | 1: 80 | Surface |
| BV421 | CD25 | 2A3 | BD | 1: 40 | Surface |
| V450 | IL-17F | O33-782 | BD | 1: 20 | Intracellular |
| BV480 | CD28 | CD28.2 | BD | 1: 40 | Surface |
| BV510 | CD27 | M-T271 | BD | 1: 20 | Surface |
| BV570 | CD45RA | HI100 | Biolegend | 1: 160 | Surface |
| BV605 | PD1 | EH12.1 | BD | 1: 20 | Surface |
| BV650 | KI67 | B56 | BD | 1: 320 | Intracellular |
| BV711 | CD69 | FN50 | BD | 1: 320 | Surface |
| BV750 | CD103 | Ber-ACT8 | BD | 1: 160 | Surface |
| BV785 | CD127 | HIL-7R-M21 | BD | 1: 10 | Surface |
| BB515 | TIM3 | 7D3 | BD | 1: 80 | Surface |
| RB545 | IFNg | B27 | BD | 1: 20 | Intracellular |
| BB630-P2 | CTLA4 | BNI3 | BD | 1: 80 | Intracellular |
| BB660-P2 | IL-13 | JES10-5A2 | BD | 1: 640 | Intracellular |
| BB700 | IL-22 | MH22B2 | BD | 1: 640 | Intracellular |
| RB744 | Granzyme B | GB11 | BD | 1: 320 | Intracellular |
| RB780 | TNFa | MAb11 | BD | 1: 40 | Intracellular |
| PE | IL1R1 | polyconal | R&D systems | 1: 20 | Surface |
| PE-CF594 | TCRgd | B1 | BD | 1: 20 | Surface |
| PE-Cy5 | CD137 | 4B4-1 | BD | 1: 20 | Surface |
| PE-Cy5.5 | FOXP3 | PCH101 | Invitrogen | 1: 20 | Intracellular |
| PE-Cy7 | AREG | AREG559 | Invitrogen | 1: 160 | Intracellular |
| PE/Fire 810 | CD40L | 24-31 | Biolegend | 1: 20 | Surface |
| Alexa Fluor 647 | Va7.2 | 3C10 | Biolegend | 1: 80 | Surface |
| R718 | IL-17A | N49-653 | BD | 1: 20 | Intracellular |
| Zombie-NIR | Live Dead Zombie NIR | NA | Biolegend | 1: 500 | Live Dead |
| APC-H7 | CD45RO | UCHL1 | BD | 1: 1280 | Surface |
| APC/Fire 810 | IL-2 | MQ1-17H12 | Biolegend | 1: 80 | Intracellular |

**B**

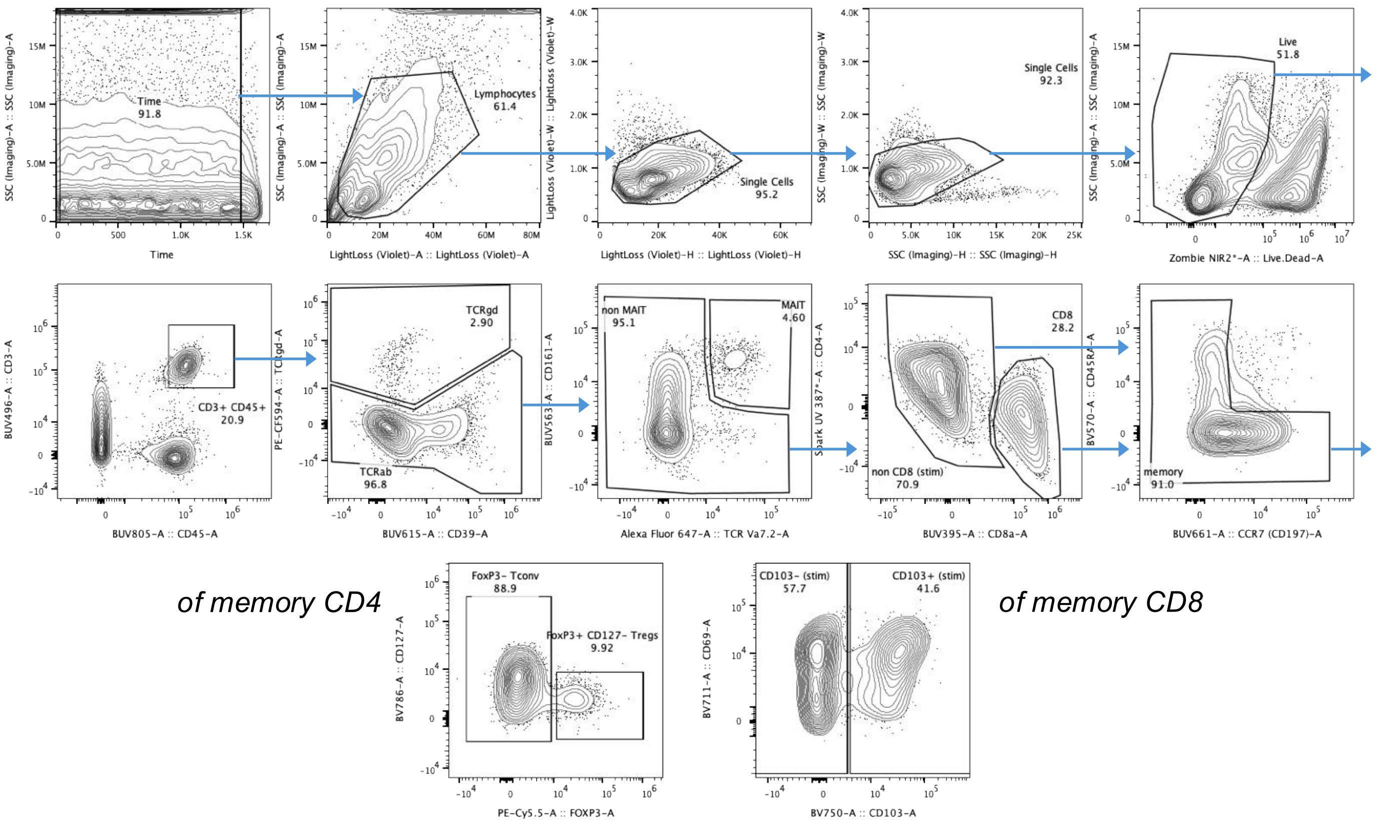

**Supplemental Figure 1: High-parameter spectral flow cytometry panel and gating scheme for analysis of tissue T cells.**

**A.** Panel information. **B.** Gating scheme: Time; lymphocytes; two single cell gates; live; T cells (CD45+ CD3+); γδ T cells (TCRγδ+) or αβ T cells (TCRγδ-); of αβ T cells, MAIT cells (CD161+ Vα7.2+) or non-MAIT; of non MAIT, CD4+ or CD8+ (CD4 is internalized upon stimulation); of either subset, memory cells (non CCR7+ CD45RA+); of CD4+ cells, Tregs (FoxP3+ CD127-) or Tconv (FoxP3-); of CD8+ cells, resident (CD103+) or circulating (CD103-). Phenotypic and functional markers assessed downstream of each subset.

### Supplemental Figure 2

A

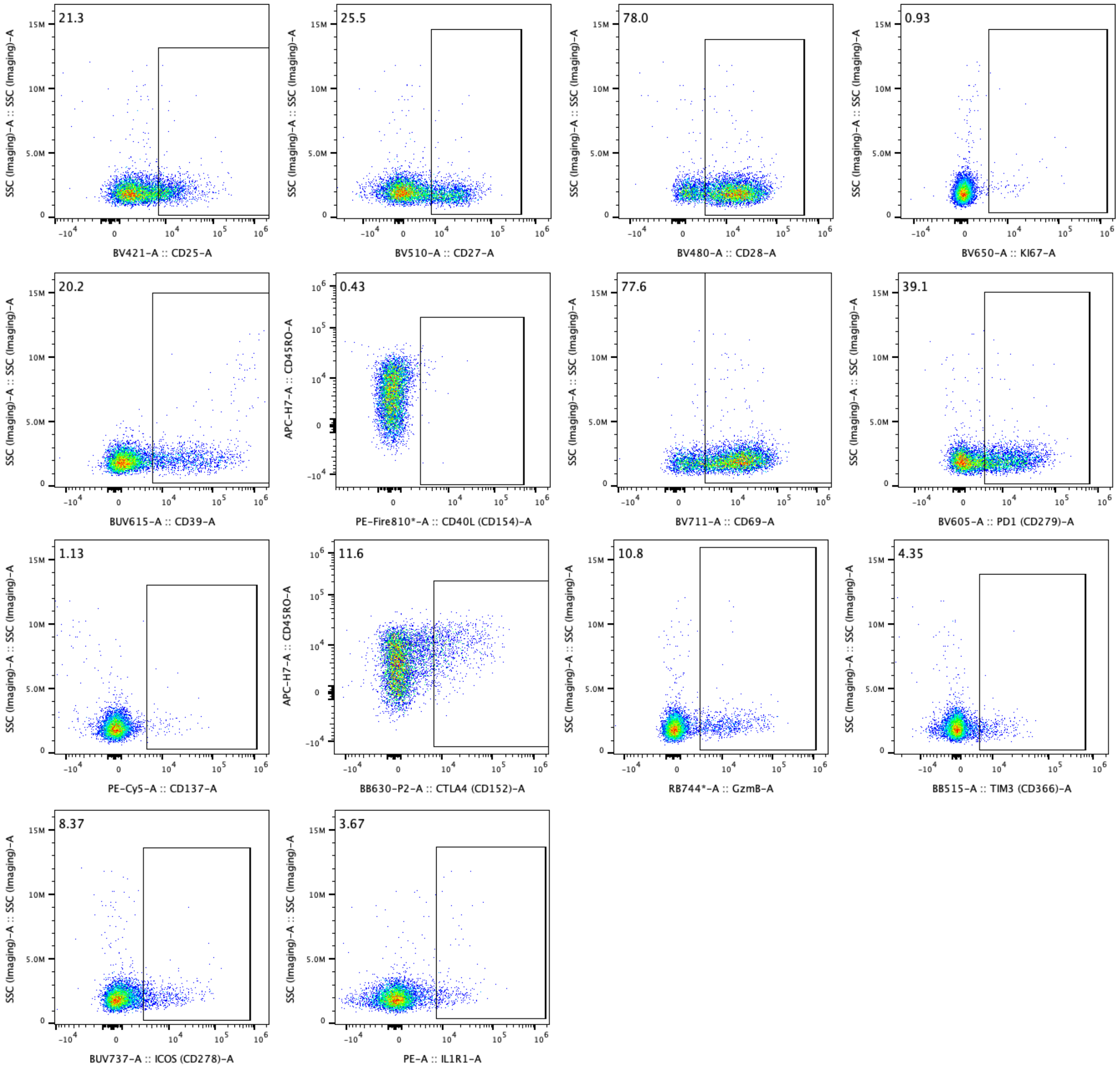


B

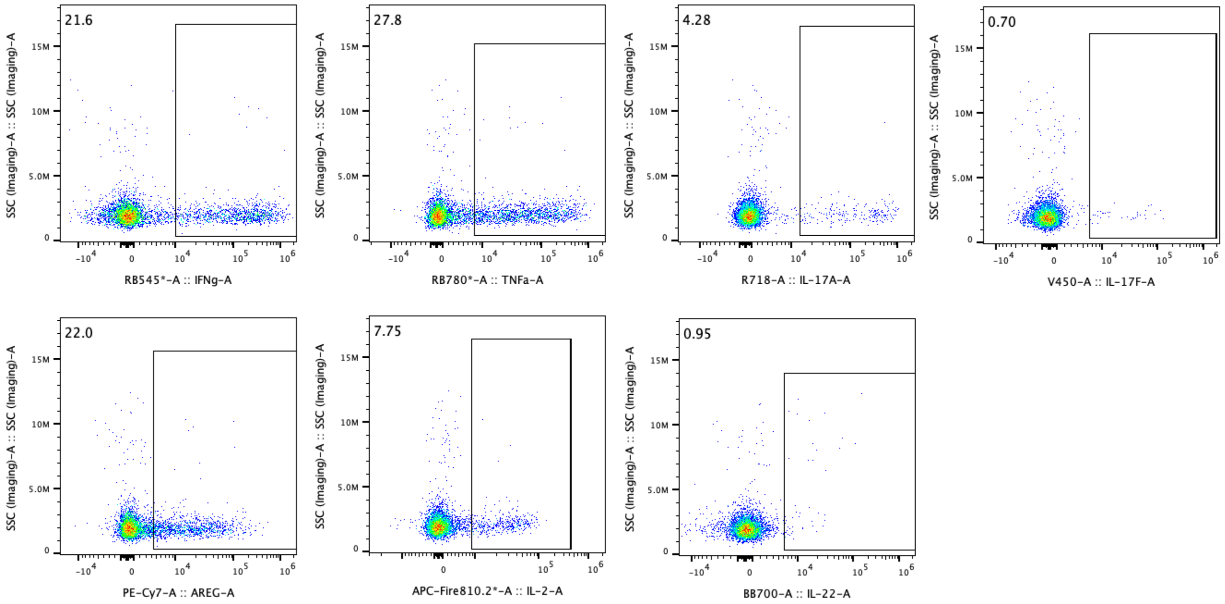


**Representative Flow Cytometry Gating**

**A.** Single positive gates of each marker pre-gated on total T cells from oral mucosa (moderately inflamed) without stimulation or **B.** upon 6-hour PMA/Ionomycin stimulation.

### Supplemental Figure 3

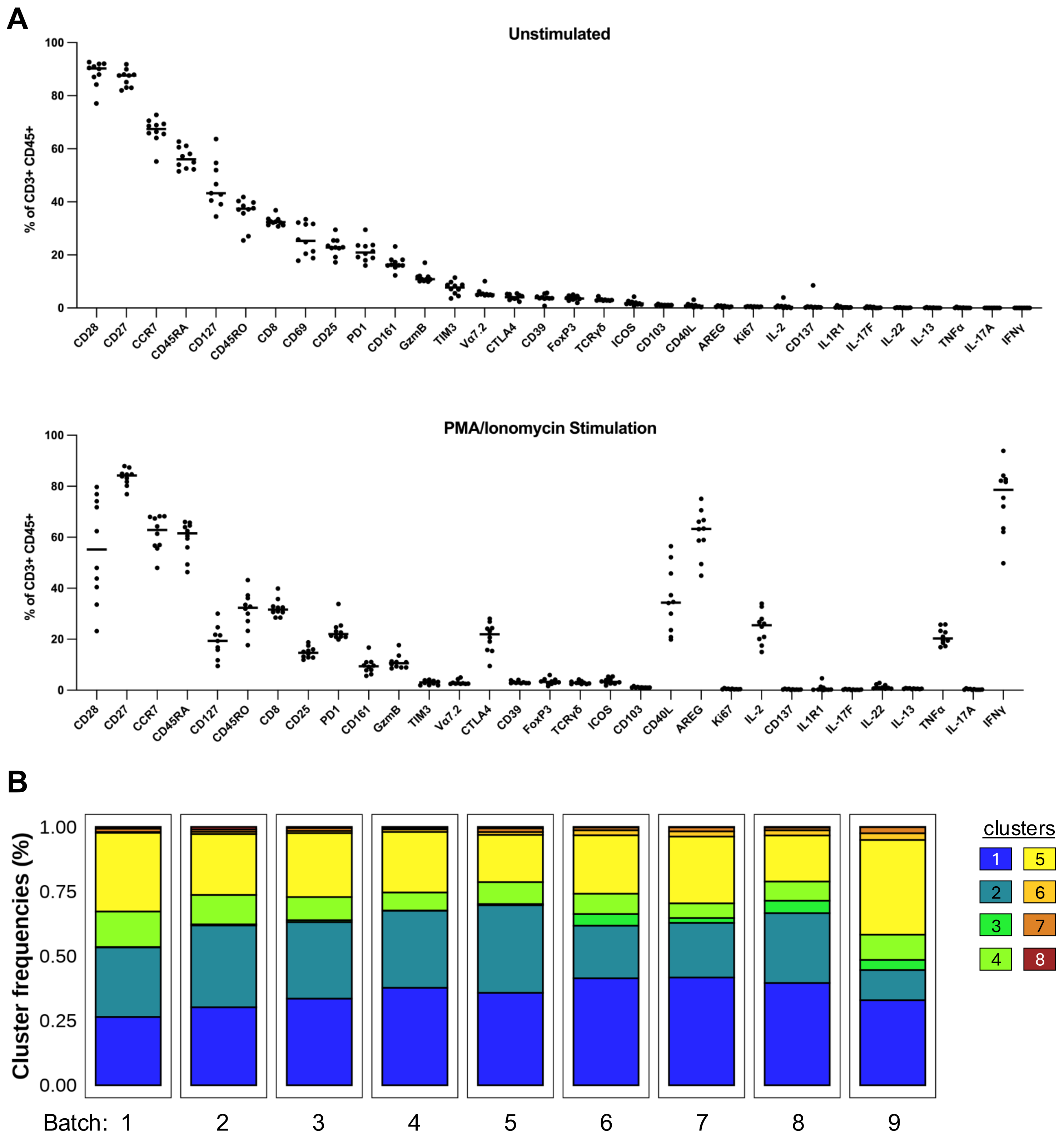


**Expression of markers *ex vivo* and upon stimulation of technical PBMC control used in flow cytometry experiments**

**A.** Consistent PBMC sample was used in every experiment to control for potential batch effects. Expression of each marker in the panel among all T cells across experiments directly *ex vivo* (top) or after 6-hour stimulation of PMA/ionomycin (bottom). **B.** Frequency of each cluster within CD4 Tconv upon stimulation (detailed further in Supplemental Figure 12) of PBMC control across experimental batches.

### Supplemental Figure 4

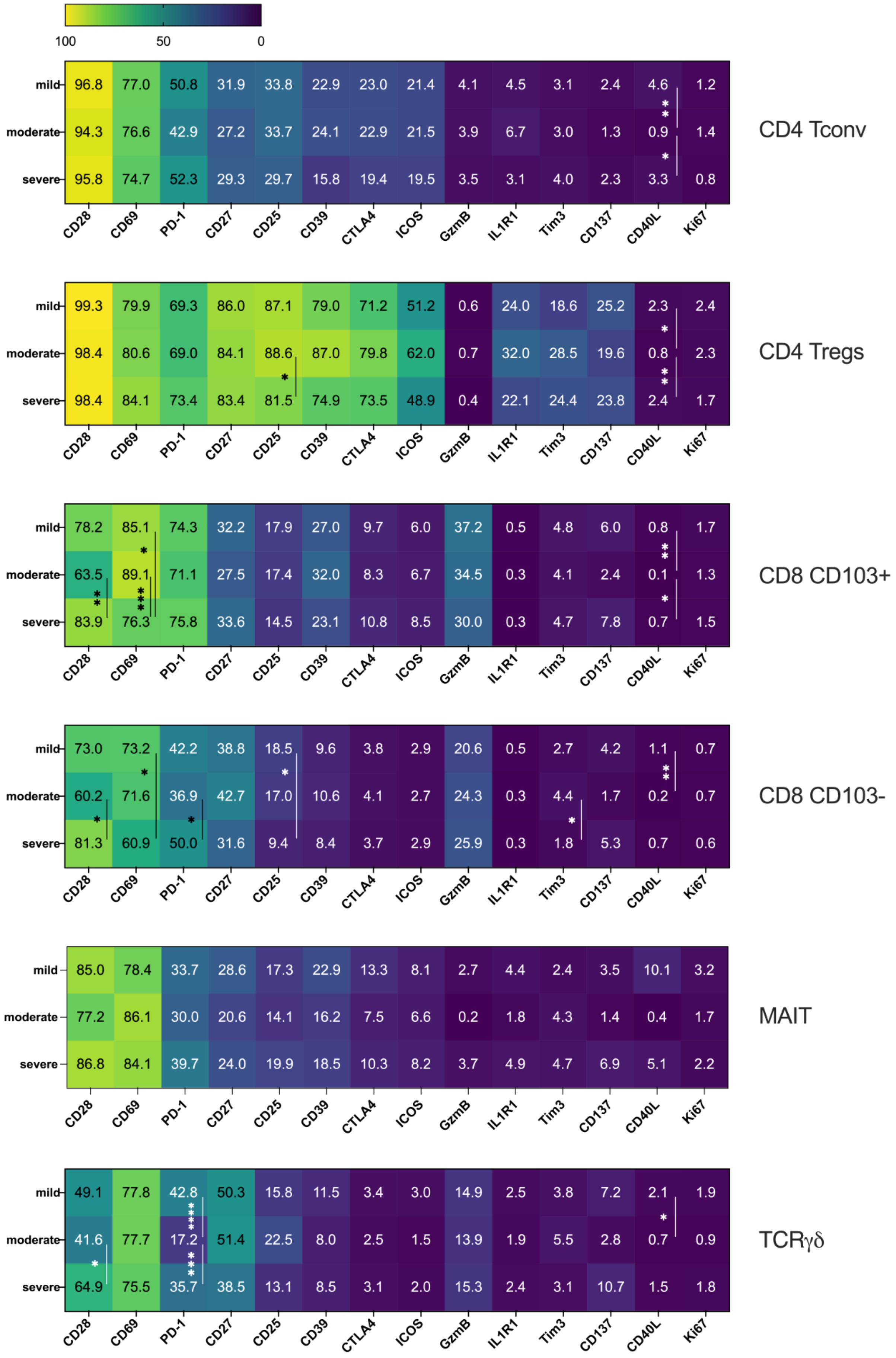


**Limited changes across inflammation status in all T cell subsets directly *ex vivo*.**

All markers assessed by spectral panel within each T cell subset in the oral mucosa across inflammation state directly *ex vivo* (no stimulation). *p < 0.05, **p < 0.01, ***p < 0.001, ****p < 0.0001 by ordinary one-way ANOVA with Tukey's multiple comparison test.

### Supplemental Figure 5

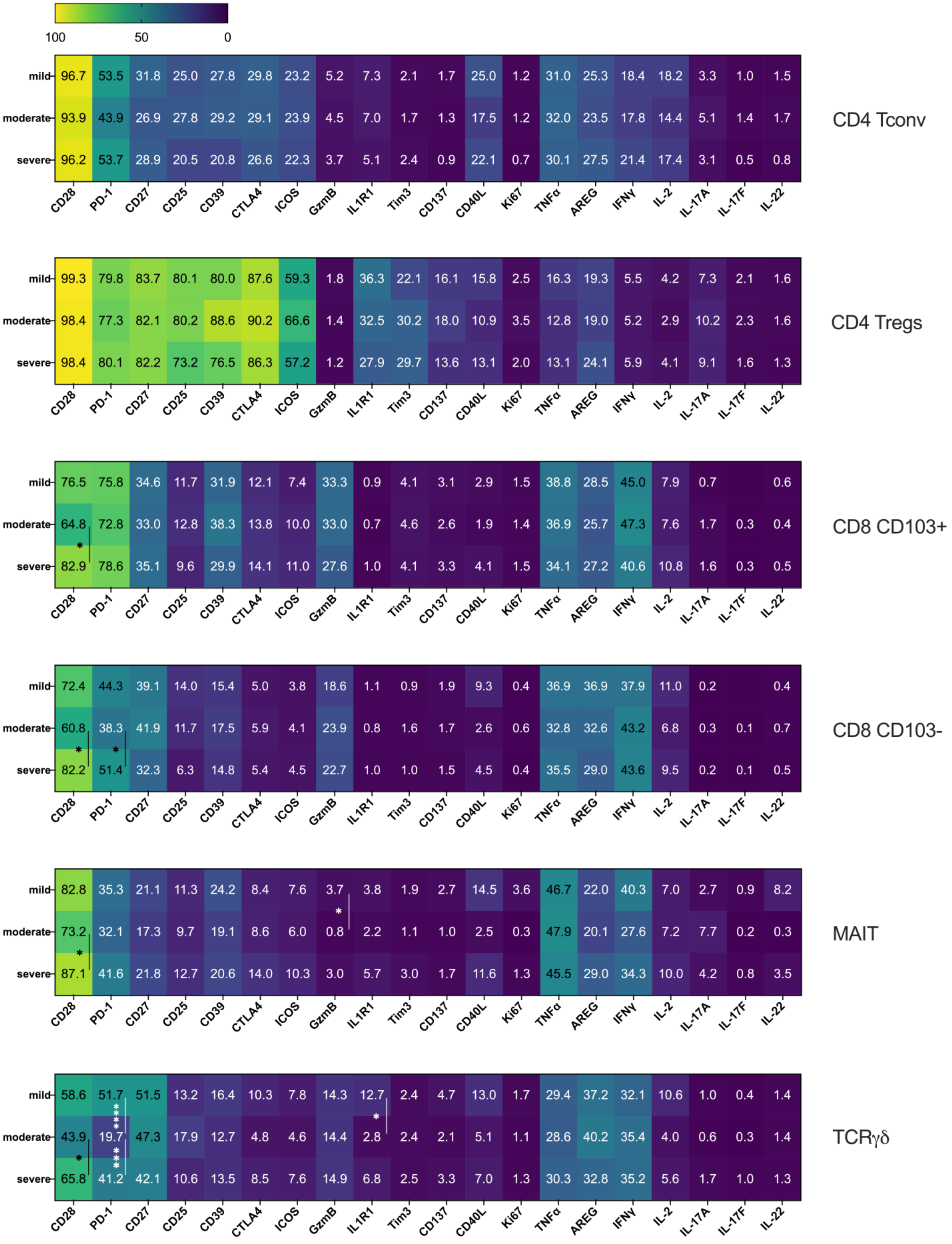


**Minimal differences in T cell cytokine production between mild, moderate, and severely inflamed oral mucosa**

Across inflammation states in the oral mucosa, all markers assessed by spectral flow cytometry following 6hr PMA/I activation stimulus. *p < 0.05, ***p < 0.001, ****p < 0.0001 by ordinary one-way ANOVA with Tukey's multiple comparison test.

### Supplemental Figure 6

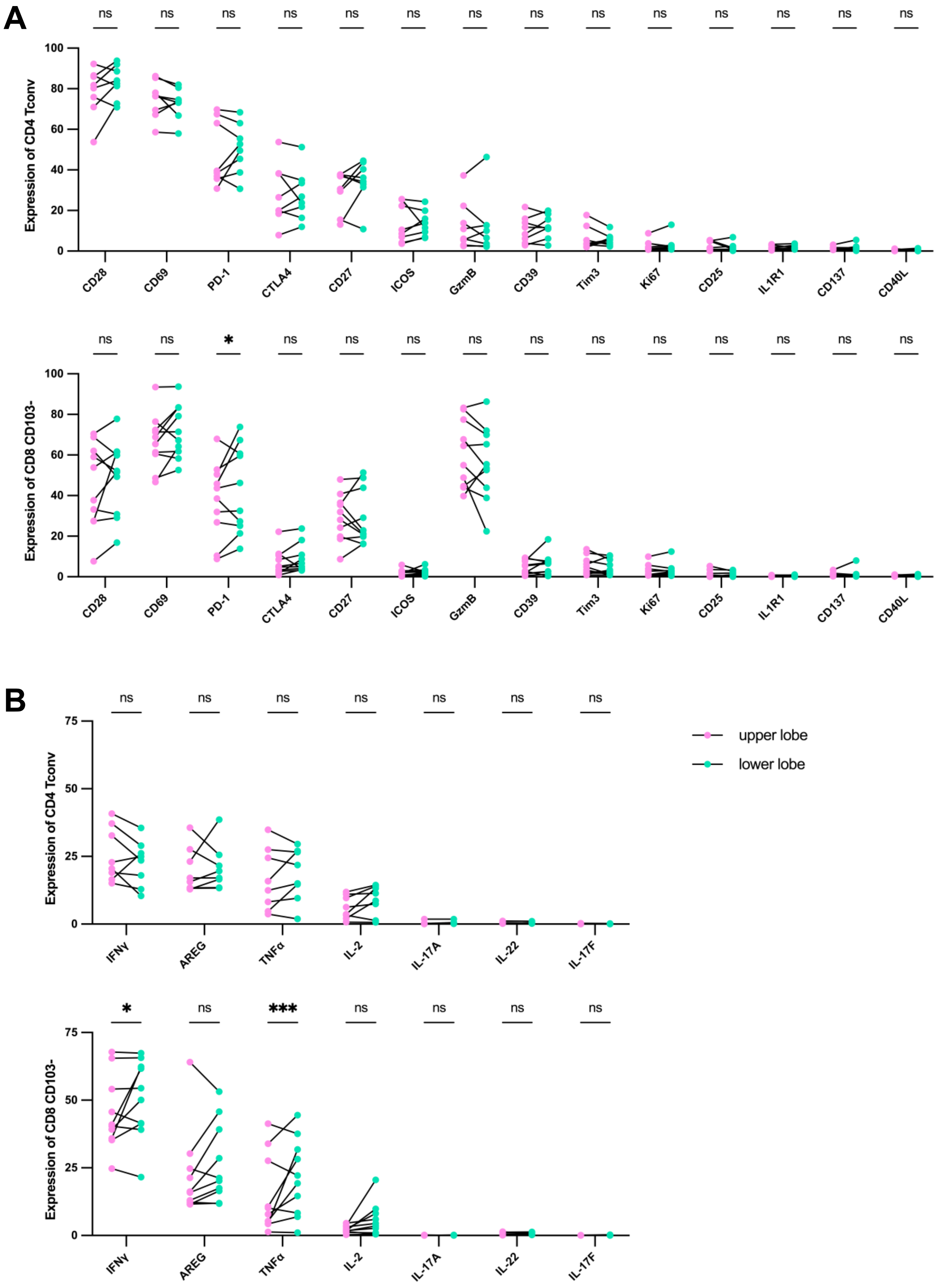


**Consistency of phenotypic and functional markers between multiple sample sites within the same donor tissue**

**A.** Frequency of each marker from unstimulated or **B.** after 6-hour stimulation of PMA/ionomycin of CD4 Tconv (top) or CD8 CD103- (bottom). Upper lobe (pink) samples are used in the remaining figures; lower lobes (green) were excluded upon determining relatively similar expression. *$p < 0.05$, ***$p < 0.001$ by 2-way ANOVA with Šidák's multiple comparison test. For one donor, two samples were taken from the upper lobe; one sample was excluded arbitrability for consistency with the rest of the analysis.

### Supplemental Figure 7

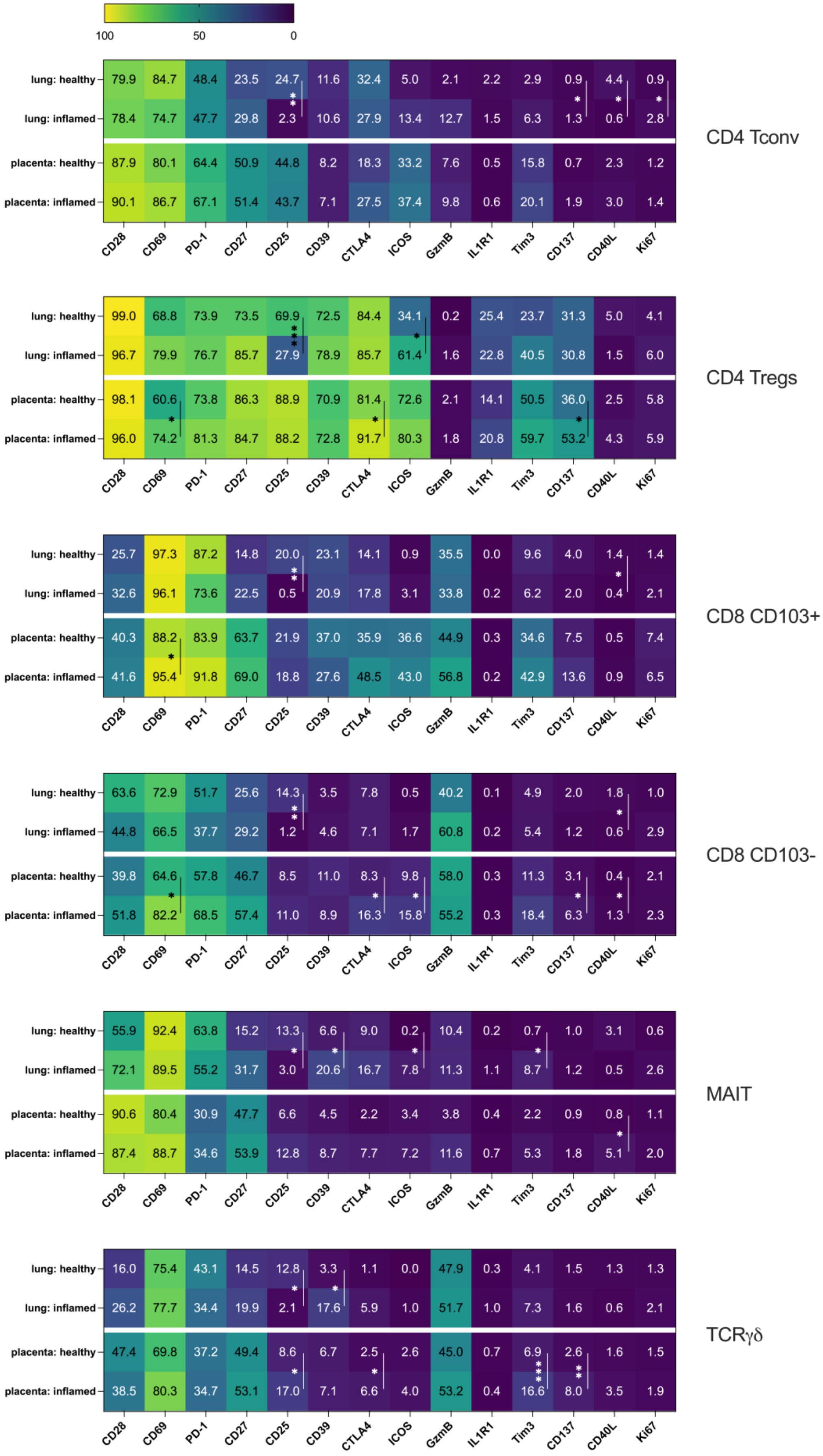


**T cell phenotype within lung and placenta tissue varies across tissue origin**

All markers assessed by spectral panel within each T cell subset in lung and placenta across inflammation state directly *ex vivo* (no stimulation). Among MAIT cells, n = 3 healthy lung and n = 9 inflamed lung. Statistical testing between tissues excluded from heatmaps. *p < 0.05, **p < 0.01, ***p < 0.001 by ordinary one-way ANOVA with Tukey's multiple comparison test.

Supplemental Figure 8

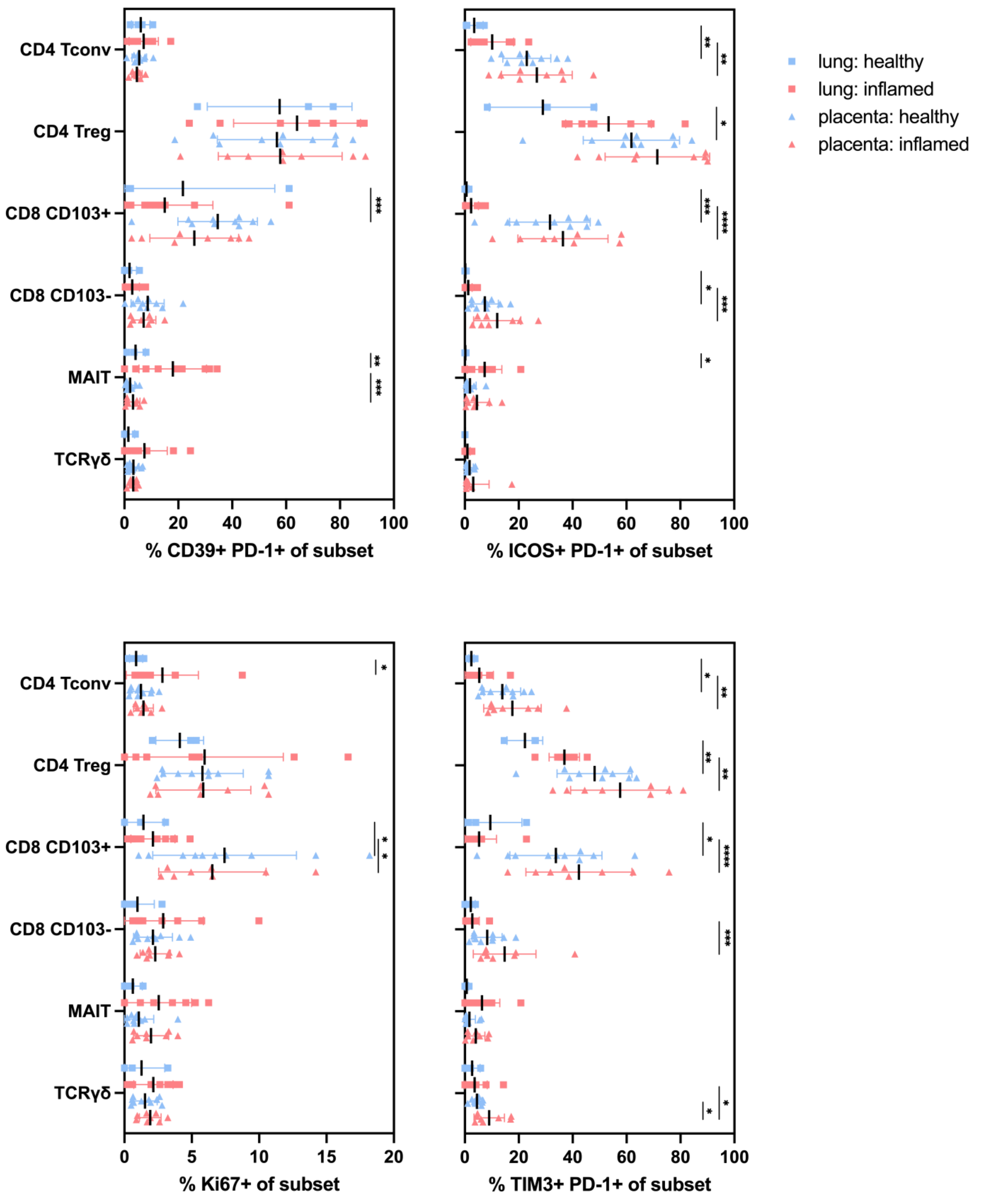


**Co-expression of biomarkers differ between lung and placenta, but are largely consistent across inflammation status**

Expression of CD39+ PD-1+ (top right), ICOS+ PD-1+ (top left), Ki67+ (bottom right), and TIM3+ PD-1+ (bottom left) among all T cell subsets across inflammation (squares, lung; triangles, placenta; blue, mild; red, severely inflamed). *$p < 0.05$, **$p < 0.01$, ***$p < 0.001$, ****$p < 0.0001$ by 2-way ANOVA with uncorrected Fisher's least significant difference test for multiple comparisons.

### Supplemental Figure 9

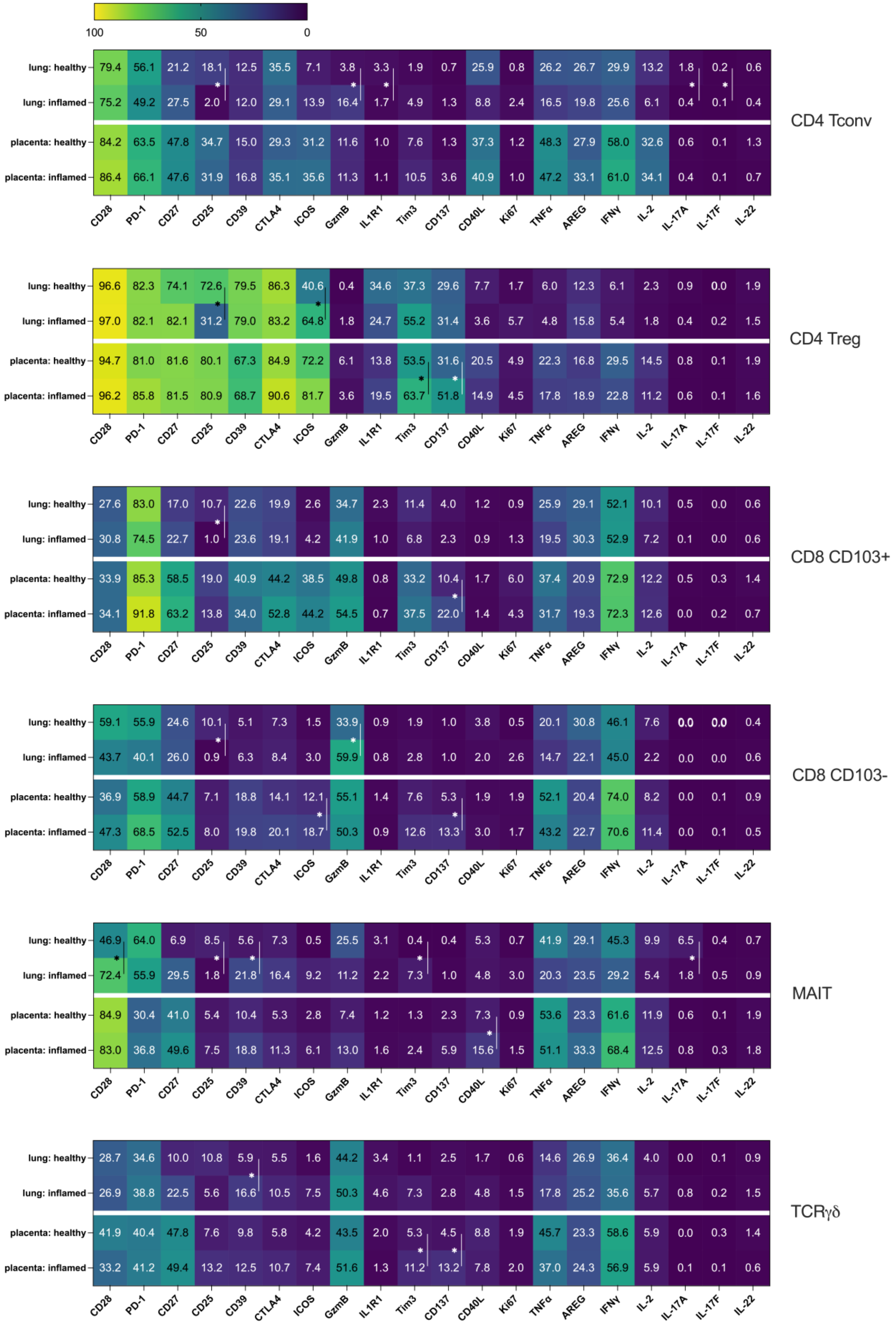

**The functional capacity of T cells, including cytokine production, are dictated largely by tissue site**

Across inflammation states in the lung and placenta, all markers assessed by spectral flow cytometry following 6hr PMA/I activation stimulus. Statistical testing between tissues excluded from heatmaps. *p < 0.05, **p < 0.01, ***p < 0.001, ****p < 0.0001 by ordinary one-way ANOVA with Tukey's multiple comparison test.

### Supplemental Figure 10

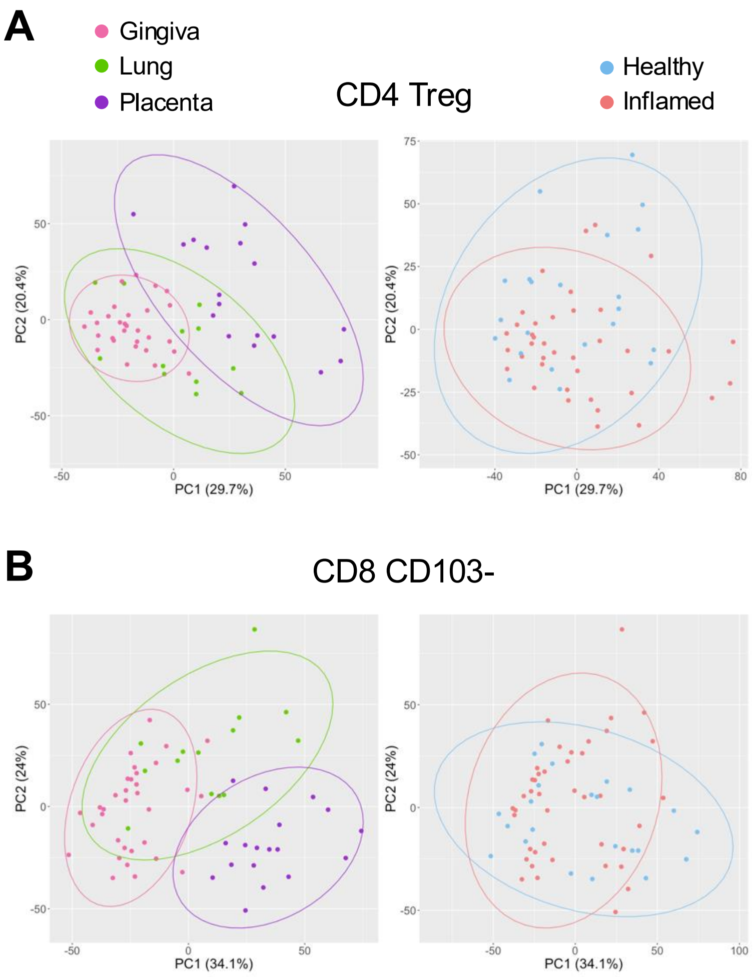


**Functional capacity of T cells groups by tissue and not inflammatory status by dimensionality reduction analysis.**

PCA plots of flow cytometry data from unstimulated samples from gingiva (pink), lung (green), and placenta (purple) during health (blue) and inflammation (red) organized by tissue type (left) and inflammation status (right) separated by subset: **A.** CD4 Treg, **B.** CD8 CD103-. For these analyses, moderately inflamed oral mucosa is grouped with inflammation category. Principal components determined by frequency of positive cells within gates from the following parameters: CD69, TIM3, CD39, ICOS, CD25, CD28, CD27, PD1, Ki67, IL1R1, CD137, CD40L, CTLA4, IL-2, IL-22, AREG, IL-17A, IFN$\gamma$, GzmB, TNF$\alpha$, IL-17F. Ellipses are 95% confidence interval.

### Supplemental Figure 11

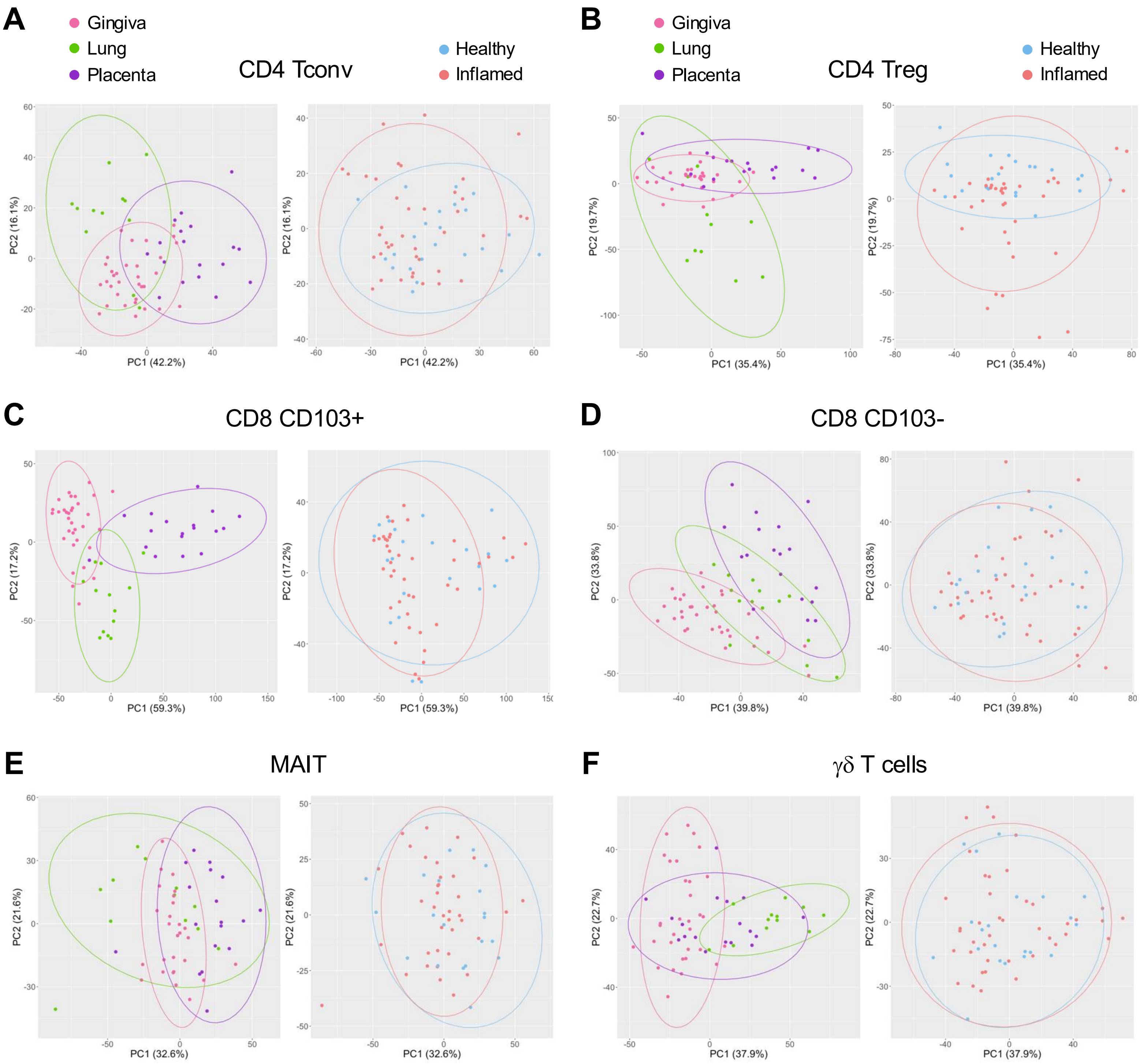


**Principal component analysis of *ex vivo* phenotype of T cells indicates separation by tissue site over inflammation status.**

PCA plots of flow cytometry data from unstimulated samples from gingiva (pink), lung (green), and placenta (purple) during health (blue) and inflammation (red) organized by tissue type (left) and inflammation status (right) separated by subset: **A.** CD4 Tconv, **B.** CD4 Treg, **C.** CD8 CD103+, **D.** CD8 CD103-, **E.** MAIT cells, **F.** γδ T cells. For these analyses, moderately inflamed oral mucosa is grouped with inflammation category. Principal components determined by frequency of positive cells within gates from the following parameters: CD69, TIM3, CD39, ICOS, CD25, CD28, CD27, PD1, Ki67, IL1R1, CD137, CD40L, CTLA4, IL-2, IL-22, AREG, IL-17A, IFNγ, GzmB, TNFα, IL-17F. Ellipses are 95% confidence interval.

### Supplemental Figure 12

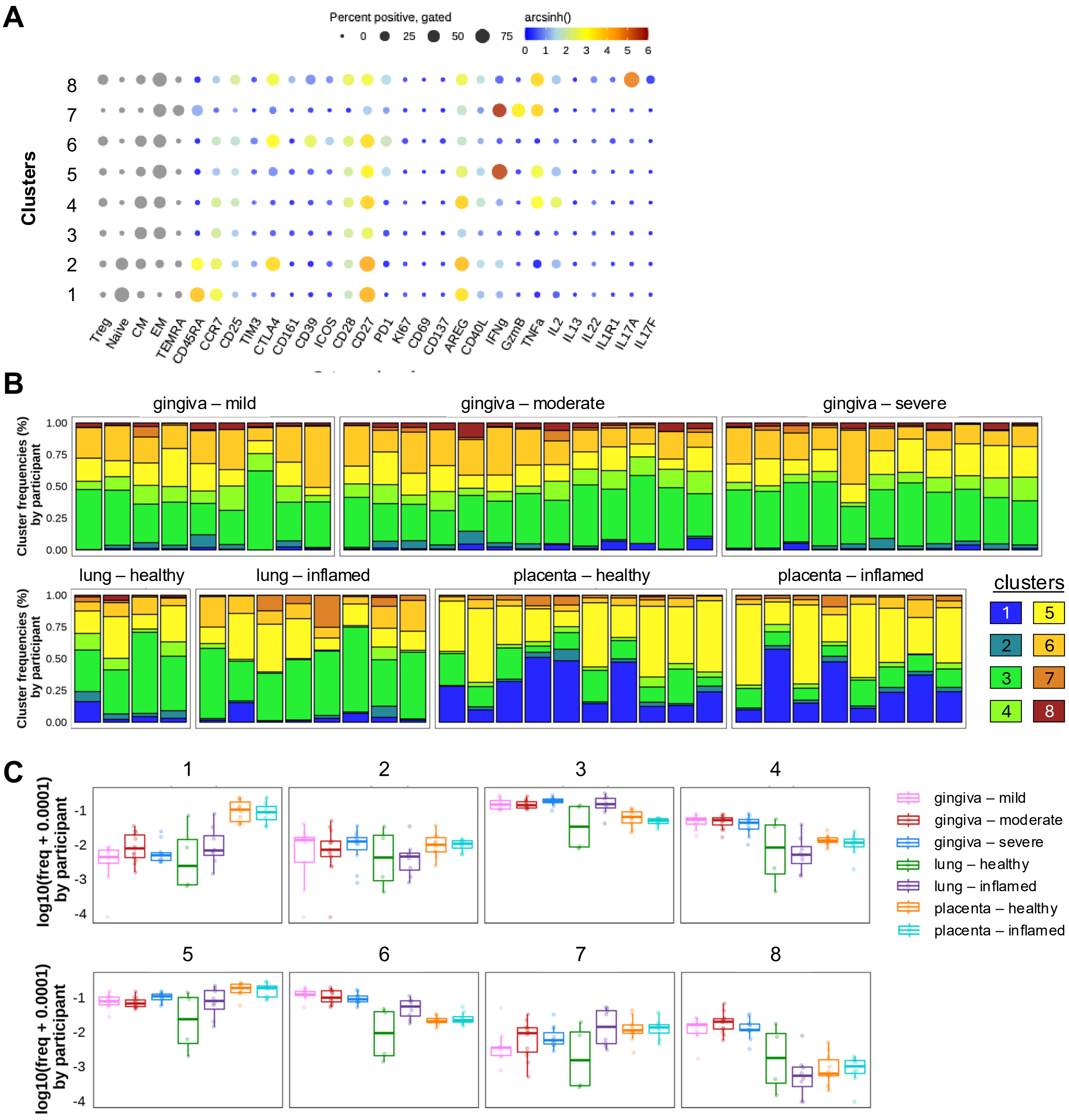


**Unsupervised clustering analysis to identify potential unique Tconv profiles based on tissue origin upon stimulation.**

**A.** Bubble plots of expression of each marker within each cluster of Tconv following PMA/I stimulation. Percentages represent overall breakdown of each cluster including all tissues analyzed (including gingiva, lung, lung associated LNs, placenta, maternal blood, tumor infiltrate and associated LNs). Resolution = 0.6. **B.** Stacked bar graphs representing cluster breakdown for individual samples. **C.** Box plots of frequency of each cluster relative to CD4 Tconv counts out of total T cells within a given sample.

### Supplemental Figure 13

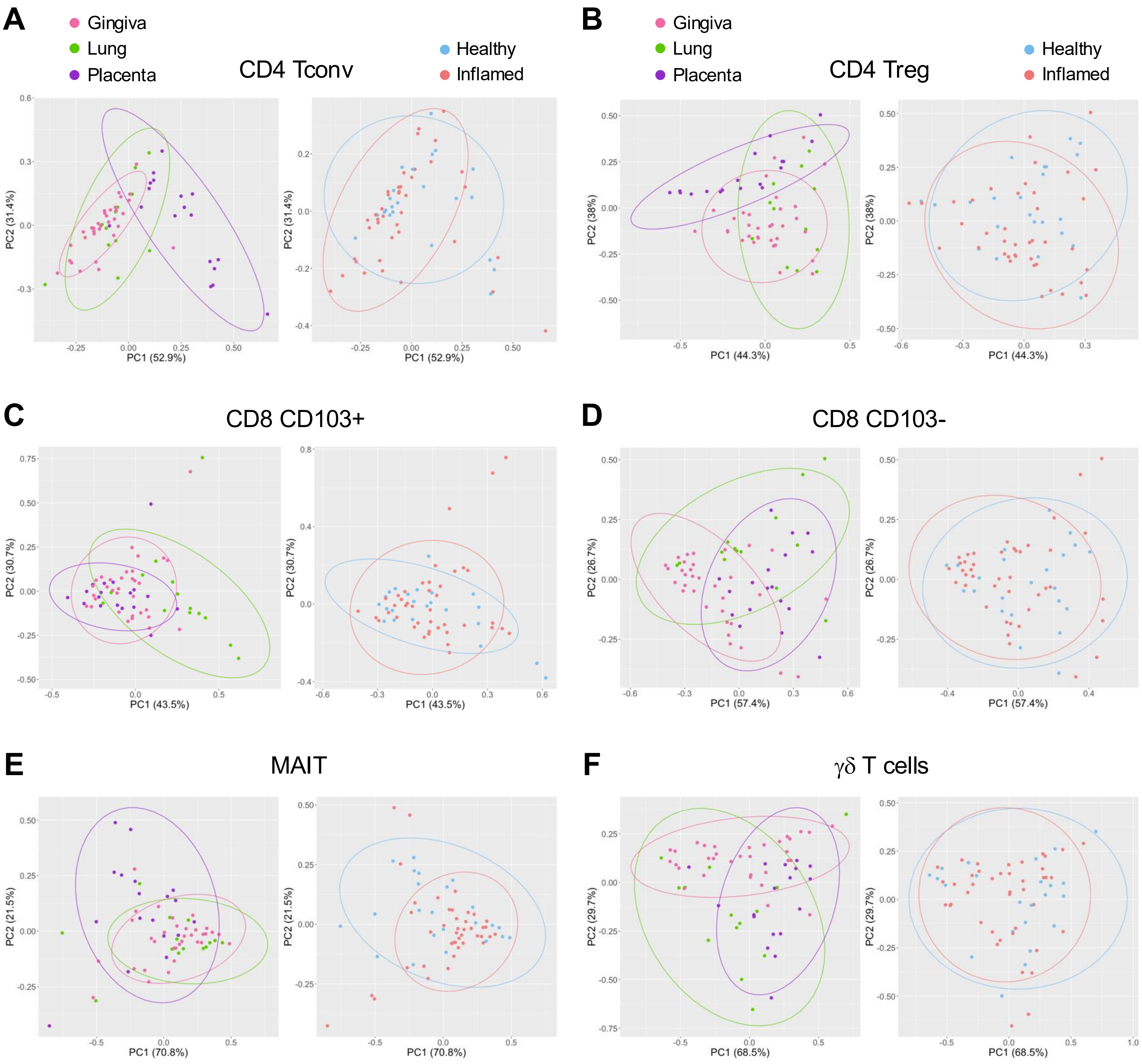


**Principal component analysis of cluster frequencies group by tissue site over inflammation status without stimulation.**

PCA plots of flow cytometry data from unstimulated samples from gingiva (pink), lung (green), and placenta (purple) during health (blue) and inflammation (red) organized by tissue type (left) and inflammation status (right) separated by subset: **A.** CD4 Tconv, **B.** CD4 Treg, **C.** CD8 CD103+, **D.** CD8 CD103-, **E.** MAIT cells, **F.** γδ T cells. For these analyses, moderately inflamed oral mucosa is grouped with inflammation category. Principal components determined by cluster frequency, described in Figures 7, Supplemental Figure 11, and data not shown. Ellipses are 95% confidence interval.

### Supplemental Figure 14

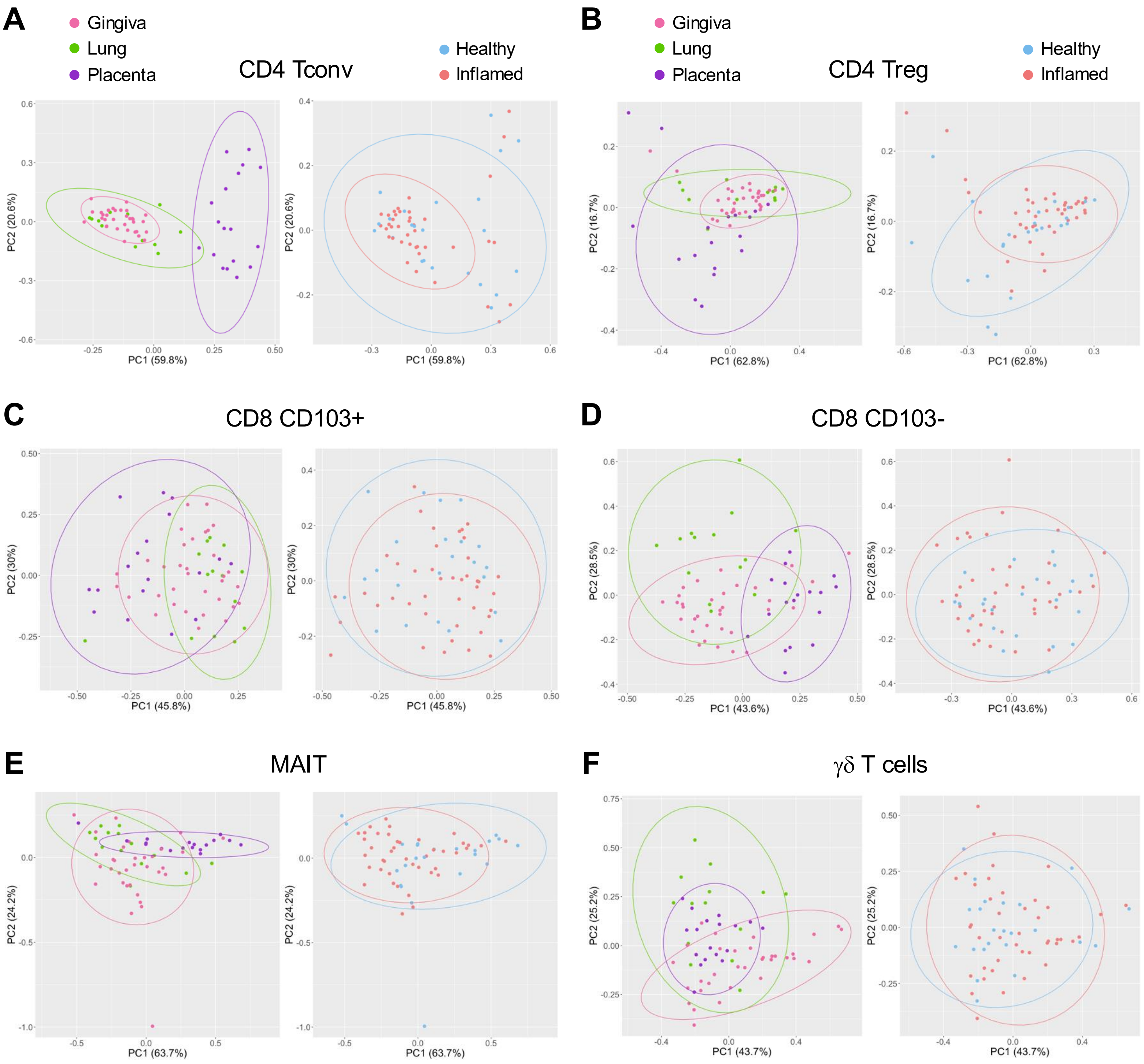


**Principal component analysis of cluster frequencies group by tissue site over inflammation status upon stimulation.**

PCA plots of flow cytometry data from unstimulated samples from gingiva (pink), lung (green), and placenta (purple) during health (blue) and inflammation (red) organized by tissue type (left) and inflammation status (right) separated by subset: **A.** CD4 Tconv, **B.** CD4 Treg, **C.** CD8 CD103+, **D.** CD8 CD103-, **E.** MAIT cells, **F.** γδ T cells. For these analyses, moderately inflamed oral mucosa is grouped with inflammation category. Principal components determined by cluster frequency, described in Figures 7, Supplemental Figure 11, and data not shown. Ellipses are 95% confidence interval.
